# Igniting full-length isoform analysis in single-cell and spatial RNA-seq data with FLAMESv2

**DOI:** 10.1101/2025.10.19.683327

**Authors:** Changqing Wang, Yair D.J. Prawer, Oliver Voogd, Jakob Schuster, Camilla Pasquali, Ricardo De Paoli-Iseppi, Anran Li, Jeannette Hallab, Luyi Tian, Hongke Peng, Margaux David, Mei R.M. Du, Silvia Velasco, Maria G. Garone, Xueyi Dong, Kathleen Zeglinski, Chiara Pavan, Kevin C.L. Law, Kwaku Dad Abu-Bonsrah, Cameron P.J. Hunt, Clare L. Parish, Quentin Gouil, Rachel Thijssen, Nadia M. Davidson, Matthew E. Ritchie, Michael B. Clark, Yupei You

**Author notes:** These authors contributed equally as co-first (Changqing Wang, Yair D.J. Prawer). These authors contributed equally as co-last authors (Matthew E. Ritchie, Michael B. Clark, Yupei You).

## Abstract

Long-read single-cell RNA-sequencing enables the profiling of RNA isoform expression and alternative splicing at single cell resolution. However, diverse single-cell technologies and sparse isoform data demand flexible and accurate analysis tools. We introduce `FLAMESv2`, a highly modular and protocol-agnostic R/Bioconductor package for long-read single-cell RNA-seq data analysis. `FLAMESv2` supports a wide range of single-cell and spatial protocols, is highly configurable, scales to allow multi-sample analysis and provides versatile visualisation and analysis outputs. We demonstrate its compatibility with both droplet-based and combinatorial barcoding single-cell methods, as well as spatial transcriptomics workflows. Benchmarking confirms `FLAMESv2` achieves field-leading performance across key analysis tasks. Applying `FLAMESv2` to *in vitro* differentiation of stem cells into neurons, we identify cell-types, differentiation trajectories, expression of annotated and novel isoforms and isoform expression diversity and heterogeneity within individual cells. `FLAMESv2` provides a comprehensive, flexible approach to analysing long-read single-cell RNA-sequencing, unlocking this powerful methodology for RNA isoform characterisation.

---

Single-cell RNA-sequencing (scRNA-seq), has become a powerful and popular method to measure gene expression within individual cells and characterise cellular heterogeneity, identify novel cell types and map developmental trajectories [1, 2]. Rapid advances in scRNA-seq technology have exponentially increased the number of cells that can be profiled, with millions of cells now able to be examined within a study [3, 4]. Such progress has driven the proliferation of scRNA-seq, including the generation of large-scale atlases of healthy, developing and diseased tissues [3, 4]. Building on scRNA-seq, spatial transcriptomics methodologies aim to profile gene expression at high resolution, enabling cellular expression profiles to be mapped within tissues. This can provide deeper insights into tissue architecture, cellular expression micro-environments and cell-cell communication [5–7]. However, the restriction of most single-cell and spatial methods to gene-level counts prevents the full-elucidation of the transcriptome.

Beyond gene-level expression, genes commonly express multiple RNA transcripts (isoforms) through processes such as alternative transcriptional initiation, splicing and transcriptional termination. Isoform expression varies between cell types and more than >90% of human genes are reported to undergo alternative splicing [8]. Alternative isoforms from a gene can have modified, or even opposing, functions and expression switches between isoforms play key roles in development, disease and cell-type specific functions [9–11]. However, while standard, short-read sequencing based single-cell and spatial transcriptomic techniques accurately quantify gene expression, they only sequence the 5’ or 3’ ends of expressed RNAs, preventing the comprehensive identification and quantification of alternative RNA isoforms. In contrast, long-read (LR) methodologies can sequence entire RNA transcripts in single reads, enabling the accurate profiling of the different RNA isoforms expressed by genes.

By combining long-read sequencing with scRNA-seq [12–17] or spatial transcriptomics [18, 19], researchers can now achieve isoform-resolved analysis at the single-cell or spatial resolution. This rapidly evolving field has spurred the development of numerous innovative sequencing protocols. However, the analysis landscape for long-read single-cell and spatial data remains highly fragmented. Many bioinformatic tools are tied to specific protocols and data types [16, 17, 19–23], with even the most flexible tools restricted to certain protocols [14, 24]. In addition, many tools require shortreads, cannot identify novel isoforms and/or do not provide an end to end pipeline and require initial data processing with another tool [20, 23–25]. These constraints limit performance, reproducibility, analytical flexibility, application to new sequencing techniques and comparisons between studies that used different methodologies. Crucially, isoform-level single-cell data is even more sparse than at the gene level, necessitating the use of accurate isoform-quantification tools specifically designed for long-read data, which have only recently become available [26]. These limitations, together with the increasing level of interest in generating these types of data, has created the need for a modular, protocol-agnostic framework that can integrate diverse data types and analysis strategies.

Here we introduce `FLAMESv2`, an enhanced and expanded `R`/Bioconductor version of our previous software [15] that offers improved flexibility, performance, speed and usability. `FLAMESv2` is compatible with a wide range of data-types, including both single-cell and spatial long-read protocols. It can be applied to data generated using long reads only, or in combination with short reads. `FLAMESv2` also provides convenient in-built functions for data visualization and analysis. Benchmarking `FLAMESv2` demonstrated it shows field-leading performance across key long-read sc/spatial-RNAseq tasks. We applied `FLAMESv2` to *in vitro* stem-cell to neuron differentiation datasets, identifying cell-type specific expression of known and novel RNA isoforms and characterising isoform diversity within individual cells. `FLAMESv2` transforms a fragmented analysis landscape into a modular, unified pipeline for isoform-level single-cell and spatial analysis.

## Results

### FLAMESv2: Methodology Overview

The `FLAMESv2` pipeline identifies and quantifies the expression of genes and isoforms from single-cell and spatial sequencing experiments. `FLAMESv2` performs six major steps: barcode demultiplexing, genome alignment, gene quantification, isoform identification, transcriptome alignment and transcript quantification (Figure 1). Briefly, sequenced reads are assigned to individual cells (and to spatial coordinates, if applicable), and mapped to the reference genome. Next, gene quantification is performed which includes UMI deduplication, a step to collapse duplicate molecules into a single read for accurate quantification. Next, `FLAMESv2` identifies isoforms and generates an experiment-specific transcriptome reference, to which the deduplicated reads are re-aligned. Finally, transcript quantification is performed to measure the expression level of each isoform in each cell. See Methods for more detailed explanation of each step.

**Fig. 1.**
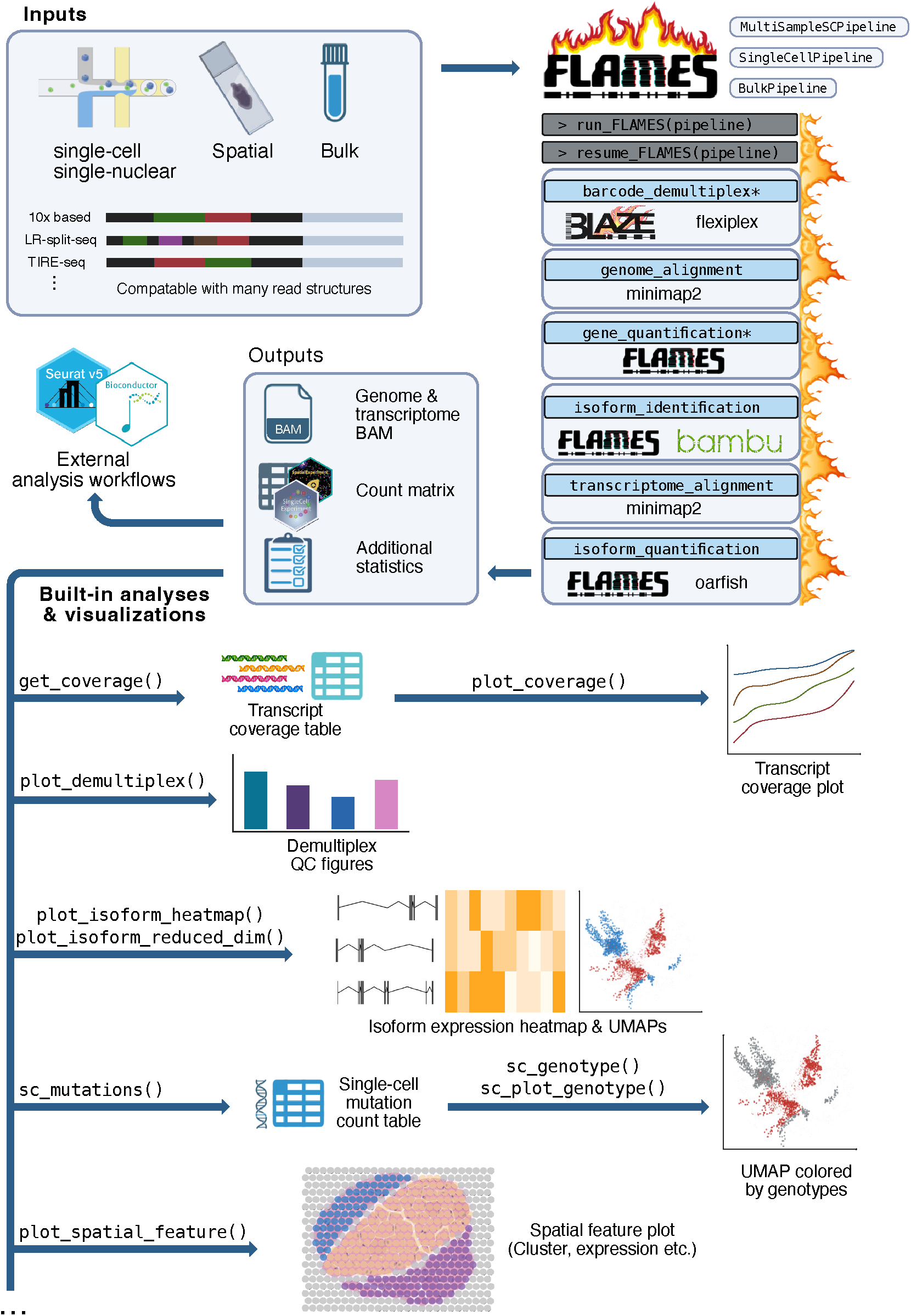
FLAMESv2 pipeline and visualization functions. `FLAMESv2` supports inputs from a wide range of experimental protocols, including single-cell (sc), single-nuclear (sn), spatial, and bulk long-read sequencing. The workflow accommodates diverse read structures and can process either individual or multiple samples. Users can perform analyses through the single-cell pipeline which is designed to handle sc, sn and spatial experiments, or through the dedicated bulk pipeline (asterisks (*) indicate steps not included in the bulk pipeline). The pipeline consists of six main stages (barcode demultiplexing, genome alignment, gene quantification, isoform identification, transcriptome alignment and isoform quantification). Outputs include genome and transcriptome BAM files, gene and isoform count matrices and summary statistics. FLAMESv2 allows flexible tool selection at key steps, enabling customization for different experimental needs. In addition to core processing, FLAMESv2 provides built-in quality control and visualization functions for data exploration and publication-ready output, including coverage plots, isoform heatmaps, genotype-annotated UMAPs, and spatial feature maps.

### FLAMESv2 provides high-performance pipelines and extensive visualization options for diverse data types

#### FLAMESv2 supports a diverse range of input data types

To maintain compatibility with the growing range of modalities amenable to long-read sequencing, `FLAMESv2` supports inputs from a wide range of protocols (Figure 1, 2A), including widely used single-cell protocols (e.g., 10x 3’ and 5’ kits, LR-Split-seq [16]), spatial transcriptomics protocols (e.g., 10x Visium), and bulk protocols, including mini-bulk protocols using well-barcodes (e.g., TIRE-seq [27]). The different protocols yield reads with different structures, hence instead of designing specific support for individual protocols, `FLAMESv2` allows users to describe their read structure via the `flexiplex` [28] package. Another major advance in `FLAMESv2`’s input flexibility is its ability to operate with or without matched short-read data. This makes the pipeline broadly accessible to researchers using only long-read sequencing, while still supporting the use of matched short-read data if available. This flexibility is achieved via the inclusion of the state-of-the art long-read demultiplexing packages `flexiplex` [28] and `BLAZE` [29] (Table S1).

**Fig. 2.**
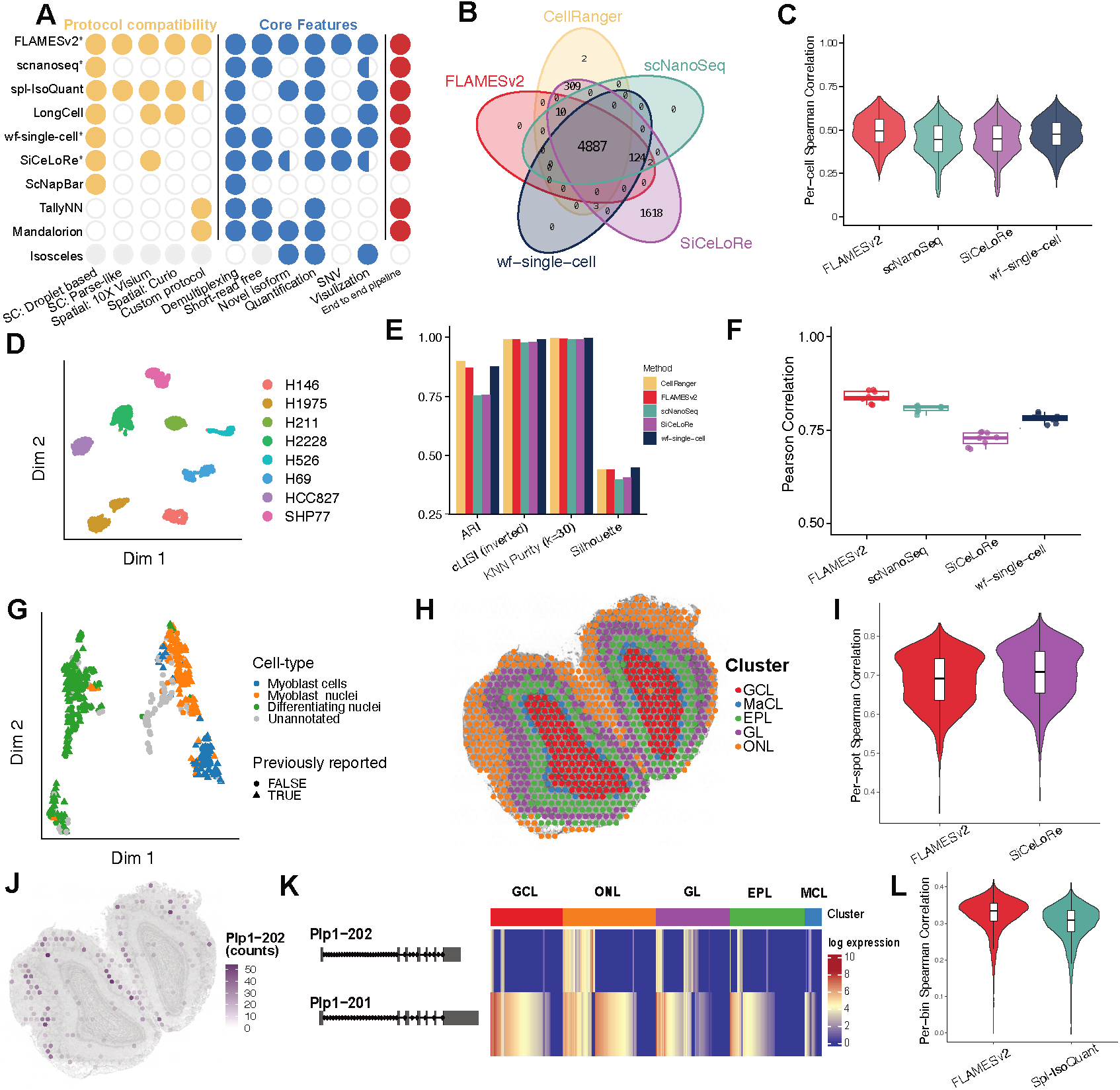
Analyzing existing datasets with performance evaluation of FLAMESv2 across various types of protocols. (**A**) Comparison of compatible protocols and core features across existing methods [14, 17, 20–25]. A filled circle indicates that the tool is compatible with the protocol or supports the feature. For visualization, a half-filled circle indicates limited visualization: software that provides only pre-configured figure generation (e.g., in a QC report) or directs users to other tools (e.g., Seurat) for analysis and visualisation. Filled grey circles indicate categories that do not apply to Isosceles, as the method assumes demultiplexed reads as input. End to end pipeline refers to methods that process FASTQ data and return single-cell count matrices. * refers to end-to-end single-cell pipelines that operate without a cell-associated barcode list from matching short-read data and instead automatically infer these barcodes from long-read data. (**B**) Overlap of identified cell barcodes across different methods using the LongBench ONT and short-read single-cell datasets. (**C**) Cell-to-cell Spearman correlation of gene-level counts between each long-read method and the short-read dataset analyzed by Cell Ranger.(**D**) UMAP visualization generated from `FLAMESv2` result on the LongBench ONT single-cell dataset, with cell lines colored according to SNP-based annotation. (**E**) Clustering performance metrics. (**F**) Transcript-level quantification comparison. Transcript counts from each method were summarized at the pseudobulk level (one pseudobulk per cell line) and correlated with bulk direct RNA (dRNA) sequencing data on the matched cell lines. (**G**) UMAP visualization of LR-Split-seq data analyzed using FLAMESv2. Previously filtered and annotated cell-type labels from short-read cells by Rebboah *et al*. were used to highlight cells and previously filtered long-read cells were denoted by point shape [16]. (**H-K**) Re-analysis and visualization of Lebrigand *et al*. (2023) [18] Visium Nanopore data with `FLAMESv2`. (**H**) Spatial plot of annotated clusters. GCL: Granule Cell Layer; MaCL: Mitral Cell Layer; EPL: Outer Plexiform Layer; GL: Glomerular Layer; ONL: Olfactory Nerve Layer. (**I**) Gene-level correlation between long-read 10x Visium data and matched short-read sequencing data from the same dataset. Long-read analysis was performed using FLAMESv2 and Scisolex (SiCeLoRe). (**J**) Spatial plot of *Plp1-202*. (**K**) Heatmap of *Plp1* isoform expression. (**L**) Re-analysis of Foord *et al*. Curio Nanopore data: Gene quantification correlation with matched short reads analysed by the vendor pipeline, comparing to the spl-IsoQuant pipeline [32]

#### FLAMESv2 offers configurable, modularised pipelines with incorporated tools

The six steps of the `FLAMESv2` pipeline are highly modular. Each step produces and accepts standard file formats, ensuring interoperability with alternative software and enabling flexible integration into diverse analysis workflows. This enables users to choose between built-in algorithms and other supported tools using the `FLAMESv2` configuration file. In addition to `BLAZE` and `flexiplex`, field-leading tools including `bambu` [30] and `oarfish` [26] can be configured for isoform identification and quantification respectively. Furthermore, many steps of the pipeline can now be selectively included or excluded, offering flexibility for different analysis needs. For example, excluding the isoform identification step will result in quantification of reference transcripts only. The modular nature of `FLAMESv2` also permits re-configuring and resuming a completed or crashed pipeline, avoiding the need to re-run jobs from scratch. To facilitate deployment on high performance computing clusters, steps can be submitted as jobs through the cluster’s job scheduler, allowing specific resource configuration for individual jobs to optimize resource usage.

#### Support for multi-sample study designs

The `FLAMESv2` multi-sample pipeline ensures uniform processing of data from each sample, including consistent naming and annotation of novel isoforms across samples.

#### Speed enhancements

Typical long-read scRNA-seq experiments now produce tens of millions to billions of reads, emphasising the importance of fast and efficient software. We have optimised `FLAMESv2` for speed. We illustrate this by processing four single-cell and single-nuclei samples from the LongBench study [31], totalling more than 300 GB of compressed `FASTQ` data, within 24 hours (Figure S1). Other core functions have been re-implemented with significantly improved speed, for example, a 10-fold improvement in single-cell mutation quantification (Figure S2).

#### Enhanced data visualization options

To allow convenient data visualization, we have added functions to create publication-ready figures. These span analysis tasks from quality control to spatial isoform expression plots (Figure 1). For example, users can process and visualize isoform expression and genotypes at specific loci on a UMAP (Figure 1). We demonstrated this function using peripheral blood mononuclear cells from a chronic lymphocytic leukemia patient [15], highlighting differential transcript usage of *RPS24* between healthy and cancer cell populations (Figure S3A) and the presence of the venetoclax resistant *BCL2 G101V* allele in a subset of cells (Figure S3B).

#### Measuring isoform diversity at single cell resolution

`FLAMESv2` includes functionality to measure isoform diversity in individual cells. The `find_diversity` function provides a direct measure of whether intra-cellular and intercellular expression is dominated by a single isoform or distributed across multiple isoforms of a gene. We demonstrate the use and biological implications of this metric in Section “Profiling within-cell diversity using Shannon’s entropy.”

#### User-focused documentation and guided tutorials

To promote accessibility, `FLAMESv2` comes with extensive user support, including detailed vignettes, tutorials and package documentation available at https://mritchielab.github.io/FLAMES, guiding users on installing and running `FLAMESv2`, using the build-in visualization functions, and performing downstream analyses.

### FLAMESv2 delivers robust, high-accuracy performance across diverse input data types

#### 10X-based long-read scRNA-seq

To assess the performance of `FLAMESv2`, we compared the pipeline against other bioinformatic tools for long-read single-cell transcriptomics (Figure 2A). While many tools address specific components of the workflow (e.g., barcode identification, isoform identification, or transcript quantification), here we focused on methods that, like `FLAMESv2`, provide end-to-end pipelines which: (i) process long-read single-cell FASTQ files and generate single-cell count matrices, (ii) do not require matched short-read sequencing, and (iii) are compatible with long-read 10x Gene Expression protocols. Accordingly, we benchmarked `FLAMESv2` against scNanoSeq [23], SiCeLoRe [14], and wf-single-cell [20].

We performed this comparison using the *LongBench* dataset, which includes Oxford Nanopore Technologies (ONT) long-read scRNA-seq data from a mixture of eight lung cancer cell lines [31]. We evaluated performance across barcode identification, quantification, and cell clustering. Matched short-read single-cell and bulk long-read datasets were used as a reference for assessing performance (Figure 2B–F, S4, S5). `FLAMESv2` identified a barcode set consistent with Cell Ranger results from matched Illumina data and comparable long-read tools, although SiCeLoRe was an outlier with many unique barcodes (Figure 2B). Similarly, UMIs and genes identified in each cell also showed strong concordance (Figure S4). `FLAMESv2` gene-level quantification showed the highest cell-cell concordance with Illumina measurements and performed on par with or better than alternative long-read methods (Spearman correlation FLAMES 0.496, SiCeLoRe 0.450, scNanoSeq 0.447, wf-single-cell 0.477) (Figure 2C).

The cell line mixture design in *LongBench* enables confident genotype-based cell annotation. All long-read tools separated cell lines effectively in low-dimensional space (Figure 2D, E, S5), yielding accurate clustering, although scNanoSeq and SiCeLoRe performed slightly worse. The *LongBench* bulk ONT direct RNA data provides an accurate reference dataset for transcript quantification [31]. We demonstrated that the aggregated pseudobulk counts per cell line from `FLAMESv2` had the highest correlation with the bulk measurements, supporting accurate transcript-level quantification with `FLAMESv2` (Figure 2F).

#### Combinatorial barcoding-based long-read scRNA-seq

To demonstrate flexible input support beyond 10x-based protocols, we reprocessed LR-split-seq data [16] (Figure S6). Four cell clusters from three samples were expected (Figure S8) [16]. As shown in the UMAP derived from `FLAMESv2`’s gene count matrix (Figure 2G), `FLAMESv2` retrieved all cells reported by long-reads in the original Rebboah *et al*. publication [16], with additional cells confirmed by the coupled filtered short-reads, and the four expected clusters showed clear separation.

Notably, the LR-split-seq custom software provides only demultiplexed reads of all possible combinations of well barcodes found in the reads, requiring manual filtering and downstream processing with additional software. To our knowledge, `FLAMESv2` is the only demonstrated method that offers a streamlined, end-to-end pipeline—from raw sequencing data to gene and transcript quantification—for Parse-like protocols. Furthermore, in addition to the barcode combination discovery mode similar to *LR-splitpipe, FLAMESv2* can also utilise the barcodes reported from the coupled short-reads and demultiplex using it as an allow-list, shown in Figure S7.

#### Spatial data

`FLAMESv2` supports long-read spatial data, such as from the 10x Visium and Curio platforms. We applied `FLAMESv2` to mouse olfactory bulb data from Lebrigand et al. (2023) [18], as well as to human visual cortex data from Foord et al. (2025) [32], generated using the 10x Visium and Curio platforms, respectively. `FLAMESv2` successfully recapitulated the reported spatial distribution of major cell types (Figure 2H) and library size (Figure S9), and achieved gene-level quantification accuracy comparable to the tools used in the original studies(mean Spearman correlation *FLAMESv2*: 0.686 and *SiCeLoRe*: 0.704 for the Visium dataset, *FLAMESv2*: 0.325 and *spl-IsoQuant*: 0.300 for the Curio dataset), despite the fact that those tools can be parameterized to optimize performance for a particular dataset (Figures 2I, L). Although transcript-level performance cannot be directly benchmarked without ground truth, `FLAMESv2` reproduced the spatially specific isoform switching of the *Plp1* gene, as visualized in heatmaps and spatial plots (Figure 2J, K), consistent with previous findings [18].

### FLAMESv2 enables benchmarking of sequencing protocols with a unified pipeline

The extensive variety of inputs, quality metrics and visualisations enables convenient technology and protocol bench-marking with `FLAMESv2`. We showcase this using the *LongBench* single-cell and single-nucleus dataset. Using `FLAMESv2`, we found the 10x 3’ single-cell kit produced more (90%) usable (demultiplexed) reads than the single-nuclei kit (64%), as shown in Figure S10 A. Of the demultiplexed reads, we observed a higher percentage with an identified TSO adapter at the 5’ end in PacBio Kinnex libraries (99%) than ONT libraries (92%), indicating more PacBio reads contain the 5’ end of the sequenced cDNA (Figure S10 B). We then compared read coverage across transcripts of different lengths for the different sequencing platforms, revealing decreased coverage and stronger 3’ bias as transcript length increased (Figure S10C). Lastly, we compared single-cell and single-nuclei isoform quantification to matched bulk RNA-seq samples. We re-processed all LongBench ONT bulk, single-cell and single-nuclei long-read samples and performed pseudo-bulk aggregation on the single-cell and single-nuclei libraries for each cell line. We then calculated their transcript count correlation with the bulk libraries. We found single-cell libraries had higher Pearson correlations with bulk data (0.72 on average) than single-nuclei (0.59 on average, Figure S10D).

### FLAMESv2 enhances isoform discovery and quantification accuracy in long-read single-cell data

To demonstrate the improved capabilities of `FLAMESv2`, we applied the single-sample pipeline to long-read scRNA-seq data generated from neuronal cultures differentiated *in vitro* from induced pluripotent stem cells (iPSCs) [33, 34]. We collected cells at day 55 of the differentiation and sequenced the sample using one PromethION flow cell, generating approximately 46 million pass reads across `∼`340 cells - (`∼`130,000 reads per cell) (Table S2). For comparison, we also processed the dataset using `FLAMESv1`.

`FLAMESv2` showed marked improvements over the original version across key metrics, identifying more unique genes (29,013 vs 5,578) and isoforms (93,721 vs 15,430), assigned a higher fraction of reads to cells (77% vs 71%) and used nearly three times more reads for isoform quantification (14.6 vs 5.1 million) (Table S2).

To assess the quality of isoform discovery, we analysed structural classifications using `SQANTI` [35, 36]. Over 95% of isoforms detected by `FLAMESv2` were classified as full-splice matches (FSM) to known isoforms. In contrast, `FLAMESv1` reported a much higher proportion of novel isoforms (45%), with only 27% classified as FSM (Figure S11). This inflated novel fraction likely reflects a higher false positive rate, driven at least in part by internal priming events that `FLAMESv1` assigns as novel due to the lack of dedicated solutions to address internal priming events. Together, these improvements reflect enhanced quantification accuracy and transcript classification in `FLAMESv2`.

### FLAMESv2 enables seamless downstream analyses

`FLAMESv2` outputs are compatible with widely used downstream analysis tools, enabling flexible and efficient exploration of long-read RNA-seq data. By supporting formats such `Seurat` [37], `Scanpy` [38] and `SingleCellExperiment` (SCE) [39] objects, `FLAMESv2` readily integrates into established workflows for gene and isoform-level expression profiling, cell-type annotation and differential transcript usage analysis. We illustrated this flexibility by analysing the day 55 neuronal culture using `Seurat` and exploring isoform-level expression across major neural populations. UMAP projections based on both gene- and isoform-level expression resolved the three major cell types expected at this stage of neural differentiation: radial glia progenitors, excitatory neurons and inhibitory neurons [33, 34] (Figure 3A). The similarity between gene- and isoform-level clustering indicates that both quantification approaches effectively capture cell-type-specific expression patterns.

**Fig. 3.**
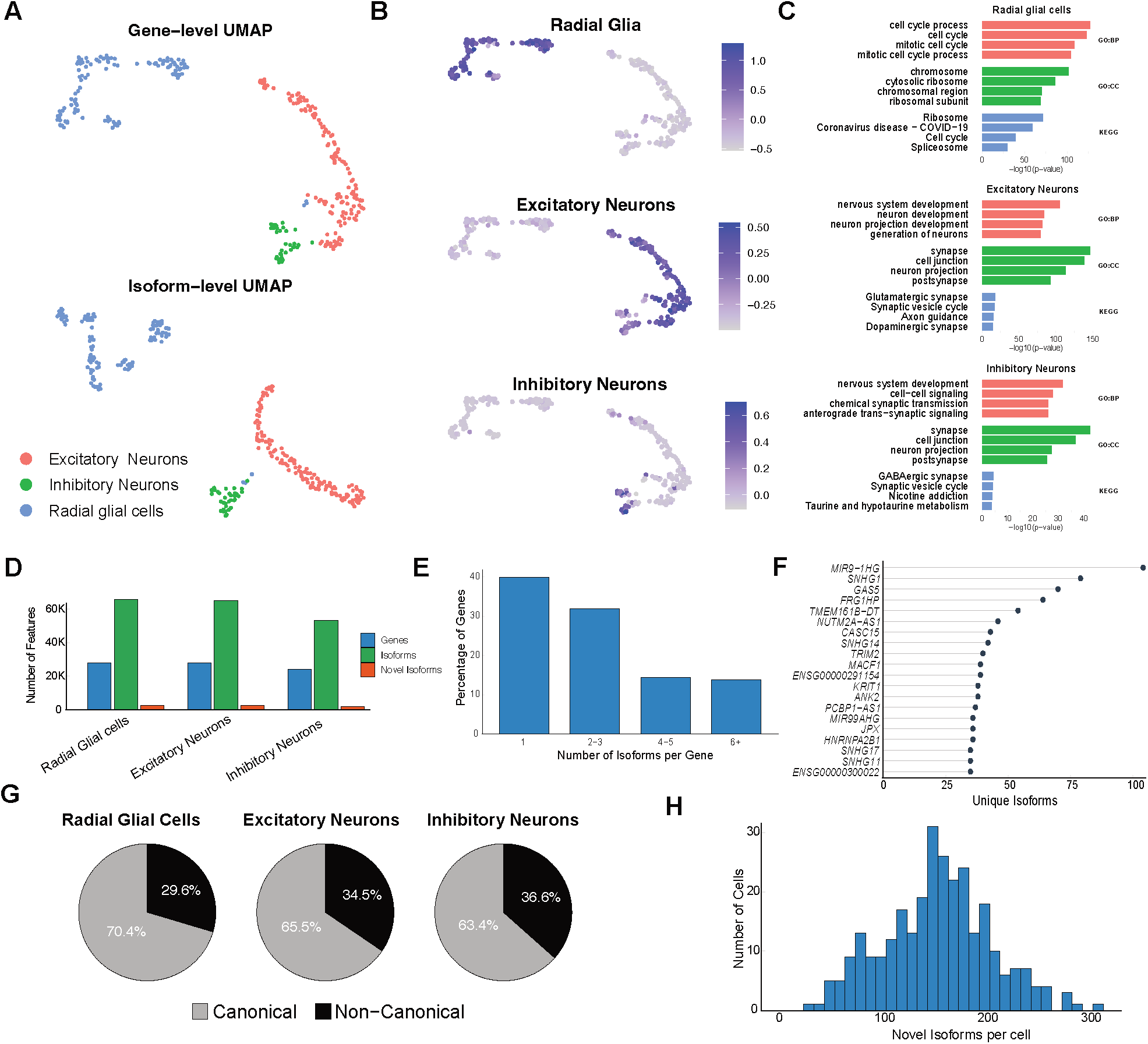
FLAMESv2 enables gene and isoform level analysis, capturing both annotated and novel isoforms across cell types. (**A**) UMAPs generated from gene-level (top) and isoform-level (bottom) expression matrices reveal three major populations: radial glial cells, excitatory neurons, and inhibitory neurons. (**B**) UMAPS overlaid with module score show relative enrichment of the selected gene set per cell and confirm cell type identities. Positive values indicate higher-than-expected expression of the gene set, while negative values indicate lower-than-expected expression. Canonical marker genes used for each gene set: radial glia (*VIM, HES1, HES5, SOX2, SLC1A3, FABP7, HOPX, NOTCH1*), excitatory neurons (*SLC17A7, SLC17A6, CAMK2A, GRIA2, TBR1, FEZF2, BCL11B, SYN1, NRGN*), and inhibitory neurons (*GAD1, GAD2, SST, PVALB, DLX1, DLX2, NPY*). **C**) Gene ontology enrichment analysis of cell-type specific genes, highlighting biological processes enriched in radial glia (cell cycle, proliferation), excitatory neurons (synapse organization, glutamatergic signalling), and Inhibitory neurons (GABAergic synapse, inhibitory signalling). (**D**) Bar plots showing the number of genes, isoforms, and novel isoforms detected across the three major cell types. (**E**) Number of identified isoforms per gene. (**F**) Top 20 genes ranked by number of expressed isoforms. (**G**) Proportion of canonical versus non-canonical isoform expression across cell types. Novel isoforms are included in the non-canonical category. (**H**) Histogram showing the distribution of novel isoforms per cell, highlighting that all cells have at least one novel isoform.

We confirmed cell identities using module scores for canonical markers (Figure 3B) and identified differentially expressed genes and isoforms with Seurat’s `FindAllMarkers`, followed by GO and KEGG pathway enrichment analysis (Figure 3C). These analyses confirmed cluster identities and uncovered biologically relevant programs, including mitotic and chromosome replication path-ways in the radial glia progenitors, as well as neuronal development and glutamatergic or GABAergic synapse formation in neuronal populations. Once cell type classification was confirmed, we explored gene and isoform diversity within each population. While the number of genes and isoforms expressed was broadly concordant across cell types, we observed fewer genes, isoforms and novel isoforms in the inhibitory neuron population (`∼`3,000, 11,000 and 750 respectively) likely a result of smaller cell numbers and reduced sensitivity to detect low-abundance features (Figure 3D). Of the 2,855 novel isoforms detected, 1,773 were ubiquitously expressed across cell types, while 176 were cell-type specific (Sup Figure S12). Over 60% of genes expressed two or more isoforms (Figure 3E), and some, such as *MIR9-1HG* and *SNHG1*, exhibited extreme complexity with over 75 unique isoforms reported (Figure 3F).

Next we explored canonical vs non-canonical isoform expression across cell types. Interestingly, radial glial cells exhibited a 5% increase in canonical isoform expression compared to neuronal populations, which may suggest that more mature neuronal cell types exhibit more diverse usage of gene isoforms (Figure 3G). Finally, we assessed the distribution of novel isoform expression across individual cells. All cells in our dataset expressed at least one novel isoform, with a near-normal distribution centred around a median of 156 novel isoforms per cell (Figure 3H). This highlights the cell-to-cell heterogeneity in novel isoform expression and the ability of `FLAMESv2` to examine lowly expressed and rare transcriptomics features. Together, these results highlight how `FLAMESv2` improves transcript discovery and quantification and provides a flexible framework for analysis.

### Profiling the isoform landscape during neuronal differentiation with the FLAMESv2 multi-sample pipeline

LR scRNA-seq tools need to be able to process growing numbers of samples and cells, enabling the analysis of RNA isoforms over development or in disease. Developing and mature brain tissues exhibit extensive alternative isoform usage [40–42], making models of neuronal development well suited to demonstrating the performance of `FLAMESv2` in investigating isoform trajectories across multiple samples and time points.

We explored isoform expression during neurogenesis using a well-established protocol for differentiating iPSCs into neurons over an 80 day time-course [33, 34] (Figure S13). We collected eight samples for single-cell processing spanning 4-time points: stem cells (D0), Day 25 (D25), Day 55 (D55) and Day 80 (D80). These time points were selected as they encompass key developmental stages, including neural induction and neuronal progenitor expansion (D25), early neuron development (D55) and neuron maturation (D80) [34]. We dissociated the cells and processed them using the 10x 3’ gene expression assay followed by sequencing on the ONT PromethION sequencer (Figure 4A), We generated `∼`949.3M pass reads (q score > 10) that were used for downstream analysis (Table S3)

**Fig. 4.**
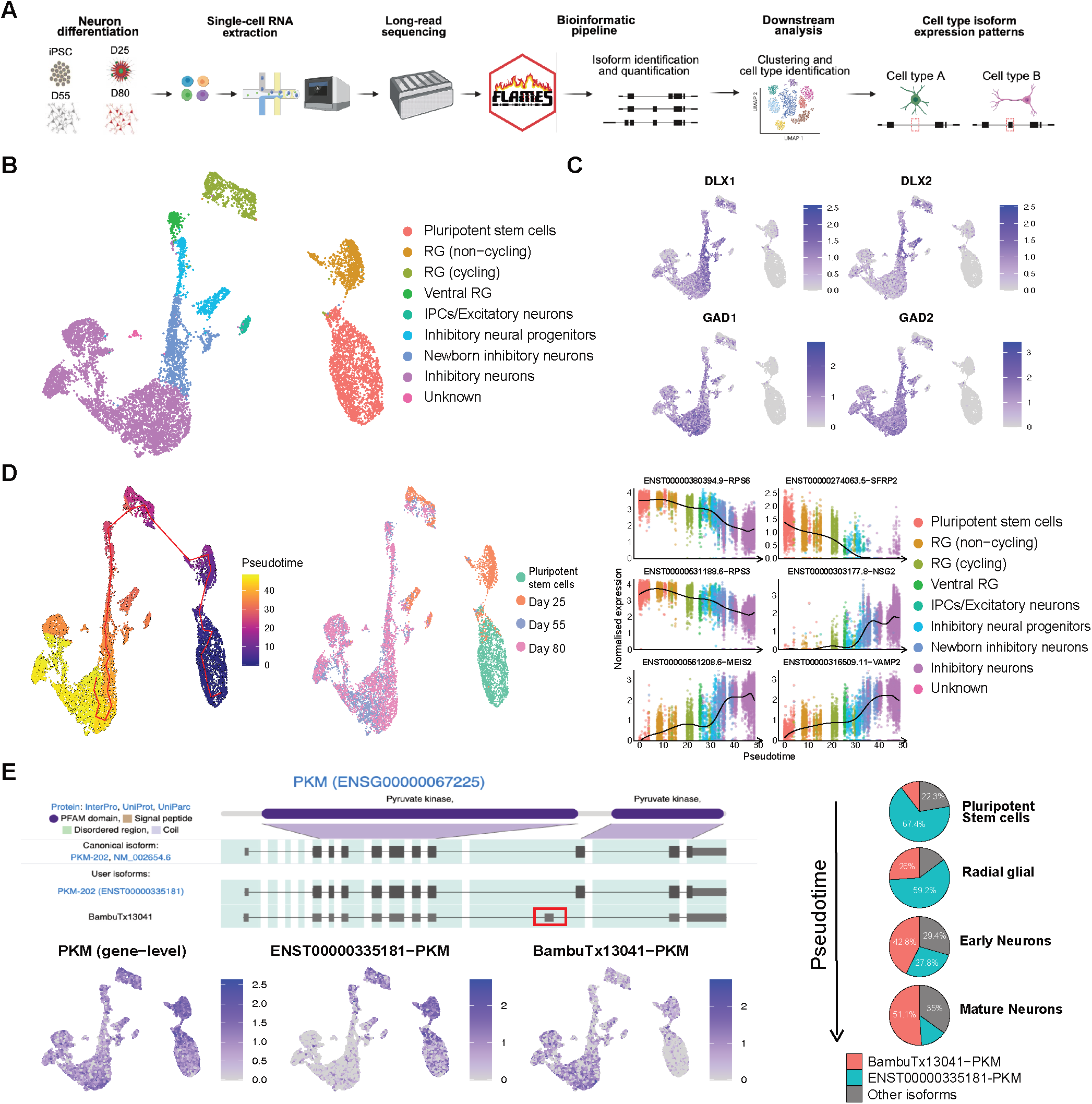
Multi sample analysis workflow in using FLAMESv2. **(A)** Experimental overview. Single-cell preparation and long-read scRNA-seq workflow using FLAMESv2. **(B)** Gene level UMAP of 8 integrated samples with annotated cell types (**RG** = Radial glia, **IPCs** = Intermediate progenitor cells). **(C)** UMAPs showing inhibitory marker expression across integrated samples. **(D)** Left: UMAP coloured by pseudotime; middle: UMAP coloured by collection day showing concordance between sampling time and inferred maturation; right: representative isoforms whose expression is strongly associated with pseudotime (three down-regulated and three up-regulated), with points coloured by cell type to illustrate the relationship between cell maturity and isoform abundance. y axis units are normalised expression values generated from `Seurat` function `LogNormalize`. **(E)** Top: IsoVis [49] image highlighting two *PKM* isoforms (canonical and a novel isoform; red box marks an exon absent from the canonical transcript). Known ORFs are denoted via thicker boxes. Bottom: UMAPs of gene-level *PKM* and isoform-specific expression demonstrate cell-type-specific isoform usage across progenitor and neuronal populations. Right: pie charts summarizing isoform proportions across broad cell classes, showing enrichment of the novel *PKM* isoform with increasing maturation.

We processed the reads using the `FLAMESv2` multi-sample pipeline, which handled all eight samples and the large number of sequencing reads in a single run. Performing isoform identification across all samples in a single pass provides greater read depth to support both known and novel transcripts, enabling improved isoform identification and quantification. In addition, multi-sample mode enables consistent isoform quantification and naming of novel isoforms across all samples, making it possible to investigate isoform expression dynamics across the time series. `FLAMESv2` identified 12,958 cells across the eight samples, 9,648 of which were retained after preprocessing in `Seurat` (see methods). We identified 1,705 to 3,099 high quality cells from each time point (Table S3, Figure S14). Across the dataset, cells contained a median of 33,993 counts assigned at the gene level and 21,272 at the isoform level. We integrated all samples using `Harmony` [43], yielding coherent batch-corrected profiles across each sample and time point.

We identified the cell types presented in the integrated dataset. As expected, at Day 0, the cultures are dominated by pluripotent stem cells. By Day 25, progenitor populations emerge (98% of cells), primarily radial glia, including a population expressing markers *HOPX, FAM107A* and *PTPRZ1* (Figure 4B, Figure S15). At Days 55 and 80, the vast majority of cells adopt neuronal identities. Following the approach in Figure 3B, we surveyed canonical intermediate progenitor cell (IPC)/excitatory neuron markers (*EOMES, TBR1, SLC17A7/VGLUT1* and *SLC17A6/VGLUT2*). Only a small cluster of 121 cells expressed these marker genes (Figure S15). Instead, expression of known inhibitory neuron markers such as *DLX1, DLX2, GAD1* and *GAD2*, showed high expression across Day 55 and Day 80 samples (Figure 4C). These data indicate that the differentiation predominately produced a population of inhibitory neurons. Previous studies have shown variability in differentiation outcomes to be a common occurrence, including the generation of more ventral or caudal inhibitory neurons, which can be driven by the cell line or batch used [44, 45]. Previously, enrichment techniques such as RNA Capture-seq have been required to properly identify such neuronal subtypes, however these were clearly defined using LR RNA-seq and `FLAMESv2` [45].

A clear developmental progression was observed along the time course. We validated these dynamics by reconstructing developmental trajectories using `Monocle3` [46] (Figure 4D, Figure S16). Pseudotime analysis recapitulated the progression from stem cells through radial glia to more mature neuronal states, consistent with known neuronal differentiation (Figure 4D). Integrating multiple samples and mapping developmental trajectories enabled us to examine both genes and isoforms whose expression changes correlate with pseudotime. We selected the 5,000 most variable genes and 10,000 most variable isoforms, then computed their correlation with pseudotime using the `Monocle3:graphtest` function. This revealed 557 genes and 216 isoforms whose expression is significantly associated with developmental progression. Notably, all of these features were also detected as differentially expressed, suggesting they capture not only discrete expression changes but also gradual, continuous shifts within the developing populations. Figure 4D highlights six isoforms whose expression was most closely correlated with pseudotime. Such patterns are particularly informative for uncovering subtle regulatory programs underlying differentiation, beyond what conventional differential expression (DE) analysis alone would reveal.

Next, we explored changes in isoform usage between cell types or states, commonly referred to as isoform switching. Using `IsoformSwitchAnalyzeR` and `DTUrtle` [47, 48], we identified 18,490, and 15,794 isoforms exhibiting significant switching between time points respectively. Of these, 11,163 were common across both methods (Table S4). Both methods identified a switching event in the gene *PKM* (*ENSG00000067225*), whose gene level expression is ubiquitous, yet shows distinct cell-type–specific isoform regulation (FDR < 0.01) (Figure 4E). Within our dataset *PKM* expresses 16 isoforms, with global expression dominated by the canonical transcript (*ENST00000335181*, 51.2% of expression) and a novel isoform (*BambuTx13041*, 31.6% of expression). The novel *PKM* isoform lacks canonical exon 9 (chr15:72202454-72202620) encoding part of the second pyruvate-kinase domain, suggesting functional divergence from the canonical transcript (Figure 4E). Isoform-level expression of *PKM* follows a clear, developmental-specific pattern in which progenitor populations are enriched for the canonical isoform, whereas neuronal populations express the novel *BambuTx13041* isoform (Figure 4E, S17). These results demonstrate that the `FLAMESv2`, multi-sample mode provides a robust framework for probing isoform-level regulation during complex processes, underscoring the package’s versatility and broad applicability.

### Profiling within cell diversity using Shannon’s entropy

Single-cell studies have traditionally focused on differences in gene or isoform expression between distinct populations (inter-cell type expression differences). Here, we leveraged our long-read data to ask two complementary questions: do individual cells express multiple isoforms of a gene (intracellular isoform diversity)? And do individual cells of the same cell type express the same set of isoforms in similar proportions, or is there appreciable isoform-level heterogeneity even among phenotypically similar cells (inter-cellular isoform heterogeneity)?

To address this, we developed the `find_diversity` function in the `FLAMESv2` package. This function calculates normalised Shannon’s entropy for each gene in each cell, providing a direct measure of whether expression is dominated by a single isoform or distributed across multiple isoforms (Figure **5**A). Because single-cell data are inherently sparse, filtering is required to ensure reliable entropy estimates. Full details of the strategy are provided in the Methods section. Importantly, all parameters are user-adjustable, allowing the function to be applied across datasets with varying sequencing depth and cell numbers.

**Fig. 5.**
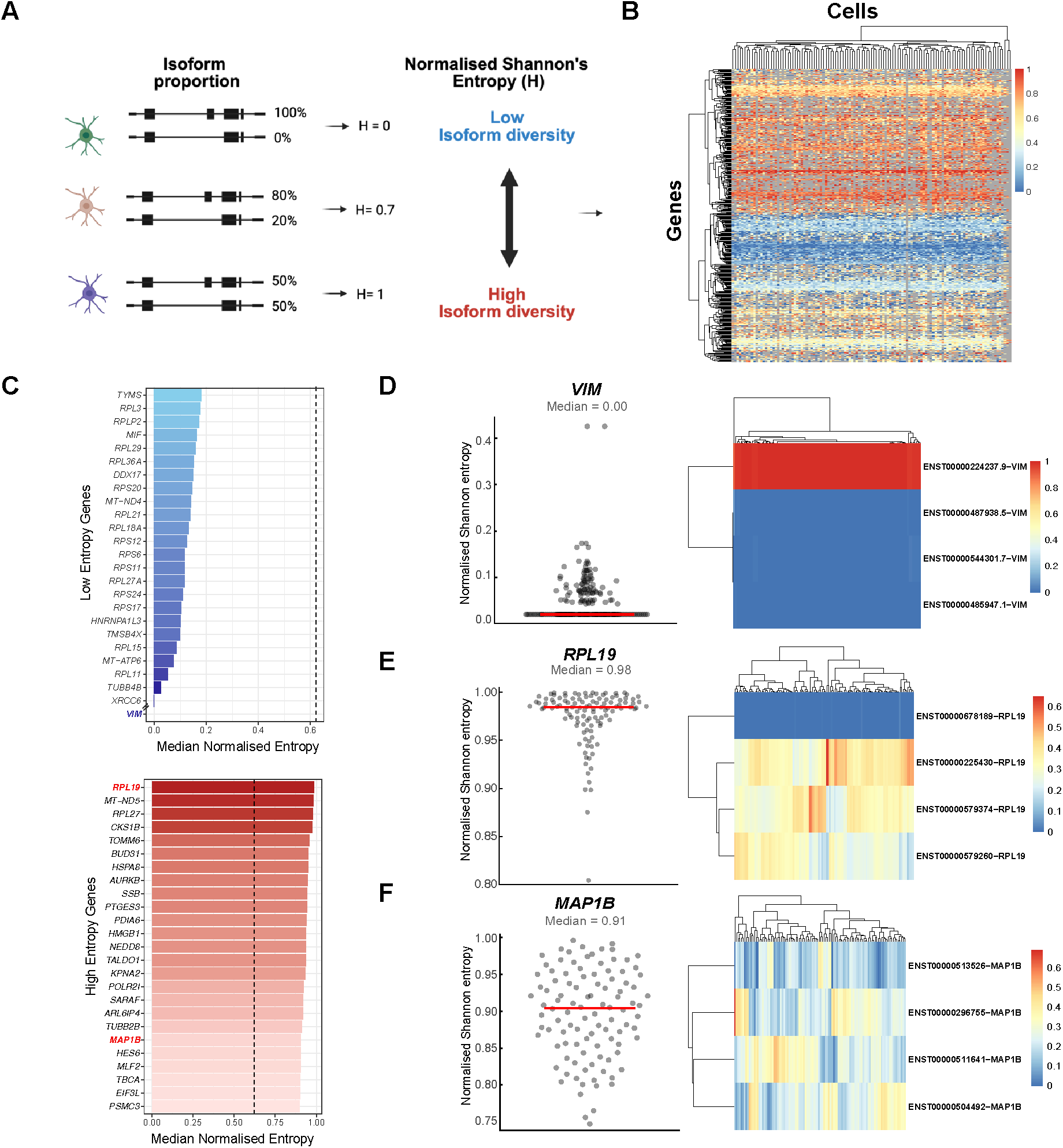
Single-cell profiling of isoform diversity using Shannon’s entropy. **(A)** Schematic illustrating how normalised Shannon’s entropy (H) captures isoform diversity. Genes dominated by a single isoform have low entropy (H *≈* 0), while those with expression more evenly distributed across multiple isoforms approach 1. **(B)** Heatmap of normalised entropy values for 435 genes across 117 radial glial cells from the Day 55 single-sample dataset. Blue indicates low normalised entropy, red indicates high normalised entropy, and grey marks cells where entropy could not be calculated for that gene due to insufficient counts. **(C)** Genes with the lowest (top) and highest (bottom) median entropy. In addition to *VIM* (shown at the bottom of the low-entropy panel), 21 genes had a median entropy of 0 but are omitted here for clarity. The dotted line in both plots indicates the median entropy values across all 435 genes. Coloured gene names represent genes shown in panels D-F. **(D-F)** Gene entropy examples. Scatter plots (left) show per-cell entropy values. Each cell is shown by a dot. Red line represents median entropy for the gene. Isoform proportion heatmap (right) shows expression proportion of each isoform in each cell. **(D)** Example of a low-diversity gene, *VIM*. **(E)** Example of a high-diversity gene, *RPL19*. **(F)** Example of a gene with heterogeneous diversity, *MAP1B*.

We applied `find_diversity` to 117 radial glial cells from our deeply sequenced Day 55 dataset (median depth: 28,540 isoform counts per cell). This approach enabled robust estimation of entropy across 435 genes. A majority of genes displayed moderate-to-high isoform diversity (24% low, 30% medium, 46% high; Figure **5**B), showing that most genes we tested are expressing multiple isoforms within individual cells (intra-cellular isoform diversity is common). Figure **5**C highlights genes spanning the spectrum of isoform diversity. Low-diversity genes such as *VIM*, which had a median entropy of 0, had nearly all expression from a single isoform (Figure **5**D). At the other extreme, highly diverse genes such as *RPL19*, (median entropy = 0.98) expressed three isoforms in all cells (Figure **5**E).

An advantage of per-cell entropy measurements is the ability to also capture heterogeneity in isoform proportions between cells of the same population. For example, *RPL19* commonly expressed isoforms in similar proportions across cells, reflecting a stable multi-isoform expression pattern (Figure **5**E). In contrast, *MAP1B* had a high median entropy (0.91) but displayed variability between individual cells. While four isoforms consistently contributed to its overall expression, their proportions differed greatly between cells, revealing inter-cellular heterogeneity in isoform composition within a cell type (Figure **5**F).

By providing gene-level diversity scores at single-cell resolution, the `find_diversity` function adds a powerful capability to the `FLAMESv2` package. This novel and complementary approach enables the investigation of isoform expression patterns between cells and states, and highlights genes with consistently high or variable diversity that may warrant further investigation.

## Discussion

Long-read scRNA-seq enables isoform-resolved analysis, revealing isoform usage across complex systems and its roles in development and disease. However, the analysis ecosystem remains fragmented. Many pipelines are tightly coupled to specific wet-lab protocols, restricting their broader applicability, while more flexible programs rely on pre-defined demultiplexing solutions designed specifically for a limited set of supported protocols, or lack desirable features such as short-read-free data processing, SNV or novel isoform identification or data visualisation (Figure 2 A).

We sought to address this issue by developing an adaptable and flexible pipeline that supports multiple data types. To achieve this, we developed `FLAMESv2` and demonstrate it can accurately process standard 10x Genomics data, as well as alternate protocols such as LR-split-seq and spatial datasets despite substantial differences in read structure. Importantly, we show that across these modalities `FLAMESv2` delivers field-leading performance in core analysis tasks, including barcode identification, gene and isoform quantification and clustering.

While `FLAMESv2` supports common analyses such as isoform identification and quantification, it is neither feasible nor desirable to hard-code every possible analysis option into the pipeline. Accordingly, `FLAMESv2` has been built to be modular, with standards-based inputs/outputs to maximize compatibility for custom steps. For example, the isoform-discovery step is tool agnostic, meaning any method that uses BAM files as input and outputs a GTF can be utilised, allowing users to swap discovery tools or benchmark them side-by-side while keeping downstream steps constant. `FLAMESv2` also integrates with the Bioconductor ecosystem and returns standard Bioconductor objects, such as `SingleCellExperiment`, for compatibility with these and other analysis packages. This design echoes other standardised and modular workflows such as `Seurat` [37]. `FLAMESv2`’s flexibility lowers the barrier to isoform-resolved biological discovery and the rich information embedded in long-read single-cell datasets. We demonstrated this potential using neuronal differentiations [33], where `FLAMESv2` quantified genes, known and novel isoforms; reconstructed developmental trajectories; and resolved cell types. `FLAMESv2` detected time-ordered isoform programs, supporting the idea that developmental shifts are also encoded at the isoform level rather than by gene abundance alone [9, 50, 51]. These isoforms provide targets for further investigation and functional follow-up. For example, we identified an isoform switch in *PKM*, where the novel *BambuTx13041* isoform encodes a distinct open reading frame that lacks part of the second pyruvate kinase domain. This domain-level rewiring likely reflects a functional shift in *PKM* function [52, 53].

Beyond differential expression, long-read scRNA-seq can quantify isoform diversity and `FLAMESv2` computes a pergene per-cell entropy score. Consistent with prior bulk studies, low-entropy genes were typically short, housekeeping loci optimized for efficient production of a single product - and high-entropy genes with richer alternative splicing profiles often related to specific developmental signatures [54, 55]. These analyses also demonstrated how individual cells can express multiple alternative isoforms of a gene and that isoform expression differs between both cells and cell types. As with any single-cell metric, data sparsity and ambiguous read alignments are potential limitations and it is recommended to restrict these analysis to genes and cells with sufficient counts for accurate isoform quantification.

Future development of `FLAMES` could further improve its performance and usability, including adding native support for empty-droplet detection and ambient RNA removal. Currently users who want decontamination must rerun parts of the pipeline with a curated background allow-list. An optional `RNA-clean` module that outputs empty-droplet/background count matrices and removes empty droplets would simplify workflows and improve quantification. In addition, while `FLAMES` is compatible with the 10x Visium spatial transcriptomics kits, future testing will be required to confirm compatibility with the recently released Visium HD platform.

The rapid advance of the long-read single-cell field also creates the need for standardised systems to benchmark experimental and computational methods. This is especially true for novel isoform discovery and quantification, as many recent studies report large numbers of novel isoforms and often infer biological importance from these findings [12, 51]. Yet, without appropriate single-cell controls, such as synthetic standards, we still have limited ability to distinguish true biology from pipeline-specific artefacts, or to quantify how accurately isoform abundances are being recovered. Within this landscape, modular pipelines such as `FLAMESv2`, where tools for individual steps can be swapped in and out, while all others are held constant, will be especially valuable.

`FLAMESv2` significantly improves on the `FLAMES` toolkit for investigating gene and isoform biology at single-cell resolution. The updated pipeline introduces substantial improvements in input support, modularity, core functionality and user experience. Users can now identify and quantify more genes and isoforms, run the pipeline with novel sequencing protocols, choose to implement the protocol with or without paired short-read data, calculate isoform diversity in single cells and visualize their data. These significant enhancements provide a faster and more versatile package dedicated to exploring genes and isoforms in single cell and spatial datasets. We anticipate that `FLAMESv2` will be instrumental in building isoform-resolved cell atlases and in pinpointing isoforms that control development and disease.

## Methods

### FLAMESv2 implementation

An overview of `FLAMESv2` pipelines are shown in Figure 1A. The first step is demultiplexing reads. The `BLAZE` and `flexiplex` packages were incorporated into `FLAMESv2` via `basilisk` [56] and `Rcpp` [57] respectively to support demultiplexing. We have updated `BLAZE` and `flexiplex` to output barcodes and Unique Molecular Identifiers (UMIs) as `HTSlib` tags in `FASTQ` files for consistency. Both packages have been modified to perform parallel `gzip` compression for the `FASTQ` output to optimize speed and storage. We modified `flexiplex` to allow specifying multiple barcodes, such as the multiple rounds of barcodes seen in Parse-like protocols [16], as well as barcodes split by inserts of fixed primers (as seen in Spl-ISO-Seq [32]). In many protocols, a sequence can be found at one end of the read to indicate it is full length, for example, the template switch oligo (TSO) from protocols based on 10x 3’ kits or the poly(dT) sequence in 5’ kits. Depending on the user’s specification, `FLAMESv2` can trim these sequences off using `cutadapt` [58]. Users may also specify to retain only reads that contain such sequences to reduce false discoveries of novel isoforms arising from truncated reads.

The demultiplexed reads are aligned to the reference genome using `minimap2` [59] and the alignment files are used for isoform identification and optional gene quantification. Next, the isoform identification step creates a transcriptome assembly, after which reads are re-aligned to this assembly using `minimap2` [59]. If novel isoform discovery is not required, users can skip this step, and the pipeline will automatically use the reference annotation instead. Isoform quantification is performed using the transcriptome alignments. `FLAMESv2` retains the option of using isoform quantification implemented in `FLAMESv1`, which is relatively conservative and assigns counts only from reads that can be confidently assigned to a single isoform, while the `Oarfish` quantification option probabilistically quantifies multi-mapping reads. Finally, a `SingleCellExperiment` object [39] containing the annotations and count matrices is returned.

#### Visualization of read coverage along transcript body

To visualize read coverage and the effect of read length on coverage, `FLAMESv2` provides a transcript coverage plot as shown in Figure 2H. For each transcript, the coverage at a specific genomic coordinate is calculated as the number of reads covering that coordinate divided by the total number of reads aligned to that transcript. Read coverage is then sampled along the transcript body, and average coverage is calculated for different transcript length ranges. To minimise the impact of misaligned reads, `FLAMESv2` uses the `filter_coverage` function to remove transcripts showing dramatic coverage change inside the transcript body (Figure S18–S19), as such patterns often reflect incorrectly mapped reads introducing artificial transcript boundaries. Specifically, the `filter_coverage` function calculates the convolution of the coverage with a step function and removes transcripts with convolutions exceeding a user-specified threshold (with the default being no more than 30% change in coverage within a sliding window of 4% of the transcript length). Coverages of individual transcripts are also available to the user to allow for further investigation with the `get_coverage` function.

### Analyses of public datasets

#### Benchmarking 10x single-cell long-read analysis tools with LongBench datasets

To benchmark 10x single-cell long-read analysis tools, we used Oxford Nanopore (ONT) single-cell sequencing data from the LongBench dataset, as most of the evaluated methods were optimized for ONT data. Because not all methods perform novel isoform discovery by default, we disabled isoform discovery in *FLAMESv2* for a fair comparison, while SiCeLoRe [14], wf-single-cell [20], and scNanoSeq [23] were run using their default settings. Additionally, ONT direct-RNA sequencing data from the LongBench project were analyzed following You et al. [31], except that we replaced the quantification tool with lr-kallisto [60] instead of oarfish [26]. This substitution was made to minimize benchmarking bias, as oarfish is the default quantifier in `FLAMESv2`. The Illumina count matrix was generated by Cell Ranger and was provided directly in You et al. [31].

#### LR-split-seq data re-processing

To demonstrate *FLAMESv2*’s capability of processing LR-split-seq data without coupled short-read data, we first processed the LR-split-seq data with *FLAMESv2*, providing the well barcode lists of the three rounds of barocdes as allow lists, for each round of barcode, *FLAMESv2* matches the candidate barcode against the corresponding barcode list, it then output reads that have all three barocdes matched within the allowed edit-distance. Using the gene counts output from *FLAMESv2*, we filtered for cells that have more than 150 UMI counts S6. Out of the 667 cells renaming, 543 were present in the filtered and annotated short-read cells from Rebboah *et al*., and we labelled the annotation in our long-read derived gene count UMAP in Figure 2G. The filtered short-read count matrices and cell-type labels were provided directly in Rebboah *et al*. [16].

#### Benchmarking long-read spatial analysis tools

We reprocessed Nanopore Visium data from Lebrigand *et al*. with *FLAMESv2* using matching genome assembly and reference annotation. We then compared the output gene count matrix from *FLAMESv2* and the SiCeLoRe gene count matrix by computing Spearman correlation against the short-read derived gene count matrix, shown in Figure 2I. The short-read count matrix and long-read SiCeLoRe gene count matrix were provided directly in Lebrigand *et al*. [18]. We also reprocessed one sample (*olderMale3*) from Foord *et al*.’s Nanopore Curio data with *FLAMESv2* and *spl-IsoQuant* version 2.2. Gene count matrix for *spl-IsoQuant* were derived using a custom script to parse *spl-IsoQuant*’s read assignment output as the pipeline does not provided single-cell/spot level count matrix by default. The corresponding short-read data were processed using Curio’s *seeker* pipeline version 3.1.0. We then aggregated the gene counts to 50x50 µm bins to reduce sparsity. Next, gene count from both *FLAMESv2* and *spl-IsoQuant* were compared against the short-read gene count matrix using Spearman correlation, shown in Figure 2L.

#### Benchmarking the Quality of Sequencing Protocols in the LongBench Project

To compare demultiplexing and read coverage between single-cell and single-nuclei samples, 10 million reads from each sample were processed with `FLAMESv2`. To identify correlations between bulk and single-cell/nuclei samples, all 24 LongBench long-read bulk samples were processed together using the `FLAMESv2` bulk pipeline, whereas all 4 long-read single-cell and single-nuclei samples were processed using the single-cell multi-sample pipeline. Alignment was performed across all 4 samples in parallel (each process was given 64 cores) as this step constitutes the primary computational bottleneck even with the highly optimised `minimap2` software [59]. For single-cell and single-nuclei samples, isoform counts were aggregated by cell-line, using LongBench’s short-read derived cell-line annotation. The aggregated counts were then used to identify expression correlations with bulk samples.

### Single-cell Sample Preparation and Sequencing

#### Human pluripotent stem cells and neuronal differentiation

Human-induced pluripotent stem cell (hiPSC) lines RM3.5 [61] and KOLF2.1J [62] were maintained under xenogeneic conditions as described by Niclis *et al*. [63]. Differentiation was performed following the protocol of Gantner *et al*. [33] as shown in Figure S13. For single-sample differentiations, RM3.5 cells were used, with culture conditions, dissociation and sequencing methods as previously described in You *et al*. (2023) [29]. For multi-sample time-course experiments, KOLF2.1J cells were used. From day 39 onwards, cultures were passaged with Accutase approximately every 10 days to reduce aggregation and ensure efficient single-cell dissociation. Apart from this modification, conditions followed those described in [33] and [29] respectively.

#### Preparation of KOLF2.1J single-cell suspensions for scRNA-seq

A total of eight samples were collected, comprising two biological replicates at each differentiation stage (Day 0, Day 25, Day 55, and Day 80). To minimize potential batch effects, differentiations were run in parallel and were staggered by approximately 30 days. This design allowed for the simultaneous collection of two samples from different time-points (e.g., Day 25 and Day 55). To disassociate cultures and prepare a single-cell suspension for 10x processing, we implemented a modified version of the “Dissociation of neuronal culture to single-cells for scRNA-seq” protocol [64]. Briefly, cultures were gently washed twice with 300 µl PBS 1x -/- to remove debris, before dissociation buffer was added. For iPSCs, the buffer contained Accutase (Thermo Fisher, Cat. #A1110501). For all other samples, the buffer comprised 2.5 ml Accutase, 2.5 ml PBS 1x, and one vial of papain (Worthington Biochemical Corporation, Cat. #LK003176). Cells were incubated at 37°C and samples were inspected every 10 minutes to ensure that breaks in the cell monolayer could be observed. Once the cell monolayer had lifted from the well, gentle trituration was performed to break apart cell clumps. The dissociation buffer was then inactivated using Wash Buffer 1 consisting of 15 mL DMEM/F12 + GlutaMAX, 15 µL ROCK inhibitor (Y-27632, 10 mM stock, Tocris Bioscience), and one vial DNase (250 µL) (Worthington Biochemical Corporation Catalog #LK003170), followed by additional gentle trituration to promote formation of a single-cell suspension. To remove any remaining cell debris the cell suspension was passed through a Flowmi™ strainer (Flowmi; Cat. No. 64709–60). Cell suspensions were washed 4 times in Wash Buffer 2 (PBS 1x plus BSA at 0.05% and Rock inhibitor (10 mM)) to remove ambient RNA contamination. Each wash was followed by centrifugation at 1200 rpm for 3 min at 4 C° with the supernatant discarded. Cells were counted manually using a hemocytometer, and viability assessed with trypan blue stain (Thermo Fisher Scientific Cat. No. 15250061). Finally, cells were resuspended in Wash Buffer 2 at a concentration of 700-1200 cells/µl as per the 10x Genomics Chromium Single-cell 3’ gene expression protocol (v3.1) user guide.

#### 10X single-cell processing and cDNA amplification

Single-cell suspensions were loaded onto the 10x Genomics Chromium Controller with a target recovery of 5,000 cells per sample, following the manufacturer’s protocol (Chromium Next GEM Single Cell 3’ Reagent Kits v3.1, CG000204 Rev D). Library preparation was carried out according to the FLT-seq protocol [65] with the reverse transcription extension time increased to 2 h to promote full-length cDNA synthesis. Following GEM generation, emulsions were split 80:20 and the 20% fraction (approximately 1,000 cells) was processed for Nanopore long-read sequencing. To minimize short fragment amplification, the cDNA was cleaned twice using 0.6× SPRI selection prior to amplification. Long-read cDNA was amplified according to the FLT-seq protocol using 12 PCR cycles followed by a final 0.6× SPRI select clean to remove primer dimer.

#### Nanopore single-cell library preparation and sequencing

Full-length cDNA generated using the FLT-seq protocol described above was prepared with the SQK-LSK114 Ligation Sequencing Kit (Oxford Nanopore Technologies, ONT) with minor modifications: incubation times for end-preparation and A-tailing were extended by 10 min, and all AMPureXP cleanup steps were performed at a 0.6× bead ratio. Libraries were sequenced on PromethION flow cells (FLO-PRO114M) with 25 fmol input. Fast5 files were generated using MinKnow v22.12.5 and basecalled with Guppy v6.3.8 (dna_r10.4.1_e8.2_400bps_sup.cfg). As per You *et al*. (2023), the Day 55 single-sample cDNA library was prepared using the SQK-LSK110 kit and sequenced on FLO-PRO002 flow cells. Fast5 files were generated using MinKnow v22.03.4 and basecalled with Guppy v6.0.7 (dna_r10.4_e8.1_sup.cfg).

### Analysing long-read scRNA-seq data of differentiating stem cells

#### Data analysis using FLAMES

To generate gene and isoform count data we used `FLAMESv2`v2.0.1 using the following configuration settings on both the Day 55 single-sample and the multi-sample time-course data. All gene biotypes - coding and non coding were quantified. Isoform discovery was performed with `Bambu` using a Novel Discovery Rate (NDR) threshold of 0.75, while isoform quantification was carried out with the default `Oarfish` model. Reads were aligned to the hg38.analysisSet reference genome with all alternative contigs removed (retaining only chromosomes 1–22, X, Y, and M). We used the GENCODE v47 comprehensive annotation [66]. Cell barcodes were identified using long-read data alone and read demultiplexing was performed with `BLAZE`. All other parameters were set to default. For all `FLAMESv1` analysis we relayed on the original implementation of read demultiplexing using a barcode allow list from matched short-reads and the original implementation of isoform discovery and quantification steps. All other `FLAMESv1` settings were the default.

#### Long-read scRNA-seq data processing and analysis using Seurat

Multi and single sample gene and isoform count matrices generated with `FLAMESv2` were processed in `Seurat` v5.1.0 [37]. The full workflow closely follows the `FLAMESv2` analysis tutorial (https://sefi196.github.io/FLAMESv2_LR_sc_tutorial/). In short, each sample was independently filtered on the gene count data to remove low-quality cells and doublets. Cells were retained if they expressed at least 3,000 genes and had mitochondrial content below 10%. We performed doublet detection using `DoubletFinder` v2.0.4 [67], setting the doublet rate = 0.039. For multi-sample analyses, gene-level data were integrated using `Harmony` v1.2.1 [43]. Isoform data was added as a new assay to the gene level Seurat object ensuring only cells that passed gene level QC were retained in the isoform assay. The isoform assay was filtered to remove isoforms expressed in less than 1% of cells. In addition, all novel Bambu genes and their corresponding isoforms were removed and not analysed in these datasets. Trajectory inference was performed with `Monocle3` v1.3.7 [46] and marker genes were identified using Seurat’s `FindAllMarkers` function with the logfc.threshold set to 0.25. Gene ontology and pathway analysis were carried out with `gprofiler2` v0.2.3 [68], using a custom background set of all genes or isoforms expressed in the filtered Seurat object. Isoforms were structurally classified with `SQANTI v5.2.1` [35, 36], with analysis restricted to those present in the filtered Seurat object (i.e., low-abundance isoforms excluded during Seurat filtering were not classified). All other parameters were kept at default unless otherwise specified. Canonical isoform classification was carried out using `BioMart` v2.6.2 [69].

#### Differential Transcript Usage

We performed differential transcript usage (DTU) analysis using `IsoformSwitchAnalyzeR v2.4.0` and `DTUrtle` v1.0.3 [47, 48]. For IsoformSwitchAnalyzeR, isoform counts were aggregated from the Seurat object using the `AggregateExpression` function, stratified by sample ID and time point (two samples per time point). Genes and isoforms were retained for testing if they reached a minimum of 30 gene level and 20 isoform level counts. Significance was defined as isoform *q*-value < 0.01 with a differential isoform fraction (dIF) cutoff of 0.25. For `DTUrtle`, we analysed the same Seurat object using the single-cell filtering option provided by the `DTUrtle` package and compared all time points (e.g. Day25 cells vs Day55 cells). Only significant hits consistently detected by both methods were considered DTU.

### Measuring cell-cell isoform variability using Shannon’s entropy

To mitigate the sparsity of single cell data, we limited our isoform variability measurement to highly expressed genes and in cells in which these genes were moderately to highly expressed. Specifically, we measured isoform variability on genes that had ≥ 100 counts and had at least two isoforms expressed in the data. For each gene, we tested isoform variability in cells with ≥ 10 counts for that gene. Isoforms were filtered to include only those expressed in *≥* 5% of cells. Additionally, within each cell, isoforms were ranked in decreasing order of abundance and filtered to include only those that cumulatively contributed up to *≥* 95% of the total gene expression. Isoforms used for variability calculation were then defined as the intersection of these *≥* 95%-cumulative isoform sets across all cells. We then used Shannon’s entropy [70] to quantify isoform variability for a given gene, similar to previous studies [71–73], but computed at the per cell level as:

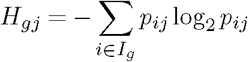

Where:

- *I*_*g*_ denotes the set of isoforms retained after filtering for a given gene *g*.
- *H*_*gj*_ is the Shannon entropy for cell *j*, quantifying isoform diversity for gene *g*.
- *p*_*ij*_ is the relative abundance of isoform *i* in cell *j*, calculated as:

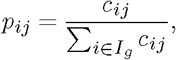

where *c*_*ij*_ is the count of isoform *i* in cell *j*.

Next, we calculate normalised entropy values 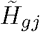 with a scale of 0-1:

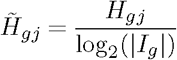

where |*I*_*g*_| denotes the number of isoforms in *I*_*g*_.

Genes were retained only if normalised entropy could be computed in at least 20% of cells. Genes were excluded from analysis if they contained isoforms with identical transcript structures (for example, that only differ in the annotated ORF). For the diversity results reported, we quantified isoforms using the `oarfish` coverage model.

## Data Availability

Cortical differentiation fastq data are publicly available to download from ENA:PRJEB100552.

## Code Availability

`FLAMESv2` is available from Bioconductor: https://bioconductor.org/packages/FLAMES. Detailed documentation and tutorials for running `FLAMESv2` and performing downstream analysis are available at https://mritchielab.github.io/FLAMES. Code for re-analyzing the public single-cell and spatial datasets is available at https://github.com/ChangqingW/FLAMES_figures, and code for data analysis of cortical differentiations is available at https://github.com/Sefi196/FLAMESv2_data_analysis.

## Author Contributions

C.W. and Y.Y. developed the `FLAMESv2` software with contributions from Y.D.J.P., O.V., J.S., L.T. and H.P. Y.D.J.P., J.H., A.L., and C.P.J.H. performed cell differentiations with assistance from C.L.P., K.C.L.L., K.D.A.-B. and C.P. Y.D.J.P. and R.D.P.-I. performed sequencing. Q.G. and R.T. generated data. Y.D.J.P., M.G.G. and S.V. performed cell type annotations. C.W., Y.D.J.P., Y.Y, C.P, M.D., M.R.M.D., X.D. and K.Z. performed data analysis. N.M.D. created research software and supervised the research. M.E.R., M.B.C. and Y.Y. planned and supervised the research. C.W., Y.D.J.P., M.E.R., M.B.C. and Y.Y. wrote the manuscript. All authors read and approved the final manuscript.

## Acknowledgements

The authors are grateful for the feedback obtained during code review of our R package when it was submitted to Bioconductor. The authors thank Dr. William Skarnes at The Jackson Laboratory for providing the KOLF2.1 cell line. This project was run as part of the 10x Genomics Millenium Science Start Single Cell Fellowship.The authors thank Dr. Catherine King and Dr. Gerry Ma (10x Genomics) and Dr. Paul Gooding (Millennium Science) for their training and assistance in the preparation of 10x libraries. The authors thank Michaela Sacco for her helpful suggestions and comments on the manuscript. This research was undertaken using the LIEF HPC-GPGPU Facility hosted at the University of Melbourne. This facility was established with the assistance of LIEF Grant LE170100200. Figure 1 and 4A was created under license with `BioRender.com`.

## Ethics approval and consent to participate

All research activities involving iPSC lines were performed under institutional ethics approval from the University of Melbourne HREC, ethics IDs: 1239208, 12374 and 28478.

