## Supplementary Materials for "Igniting full-length isoform analysis in single-cell and spatial RNA-seq data with FLAMESv2"

| Feature | FLAMESv1 | FLAMESv2 |
|---|---|---|
| Input support | 10x 3' protocol only | Extended protocol support |
| Long read only support | ✗ | ✓ |
| Putative barcode calling | ✗ (matching short-read required) | ✓ (via BLAZE or flexiplex) |
| Multi-sample support | ✗ | ✓ |
| Gene-level quantification | ✗ | ✓ |
| Novel isoform discovery | Built-in method only | Built-in method / Bambu |
| Isoform-level quantification | Built-in counting method only | Built-in counting / oarfish EM estimation |
| Modular pipeline structure | ✗ | ✓ |
| Visualizations | ✗ | ✓ |

**Table S1. Comparison of FLAMESv1 and FLAMESv2.** ✓ indicates supported features; ✗ indicates not supported. FLAMESv2 incorporates support for isoform-level quantification, modular architecture, and multi-sample workflows, enhancing scalability and interpretability of long-read scRNA-seq data.

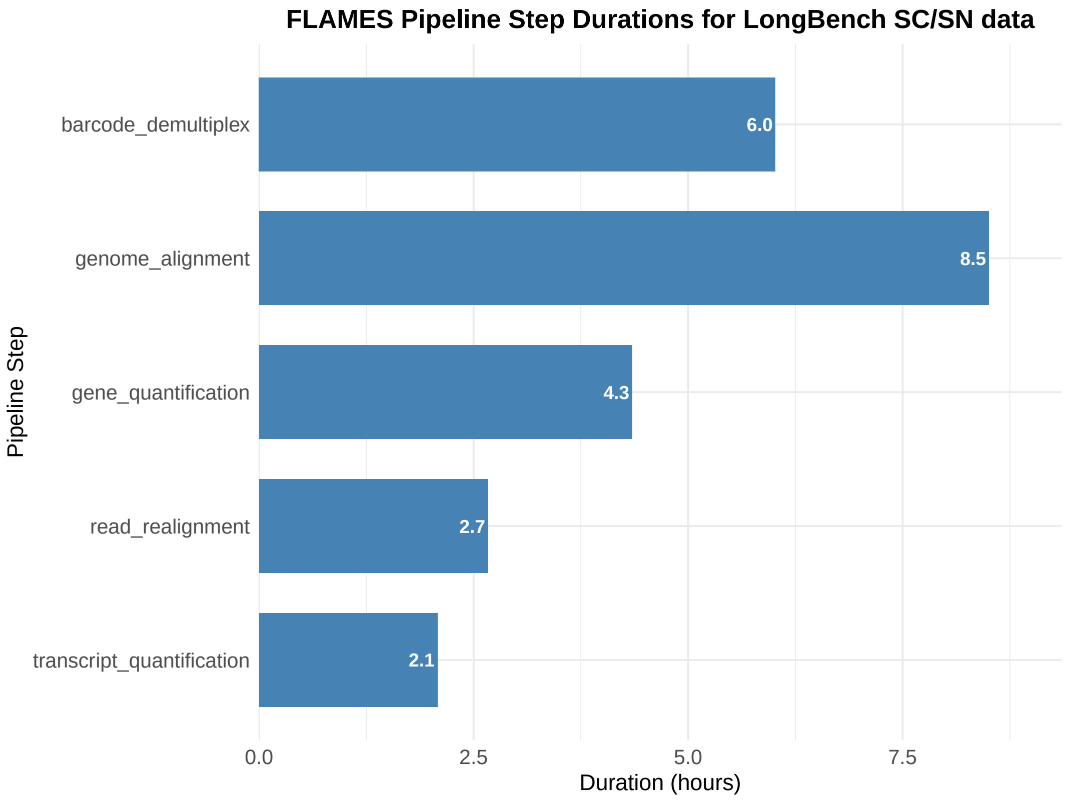


**Fig. S1. Time taken for each step in `FLAMESv2` to process all single-cell and single-nuclei data from *LongBench* [1].** Produced using the `plot_durations` function from `FLAMESv2`.

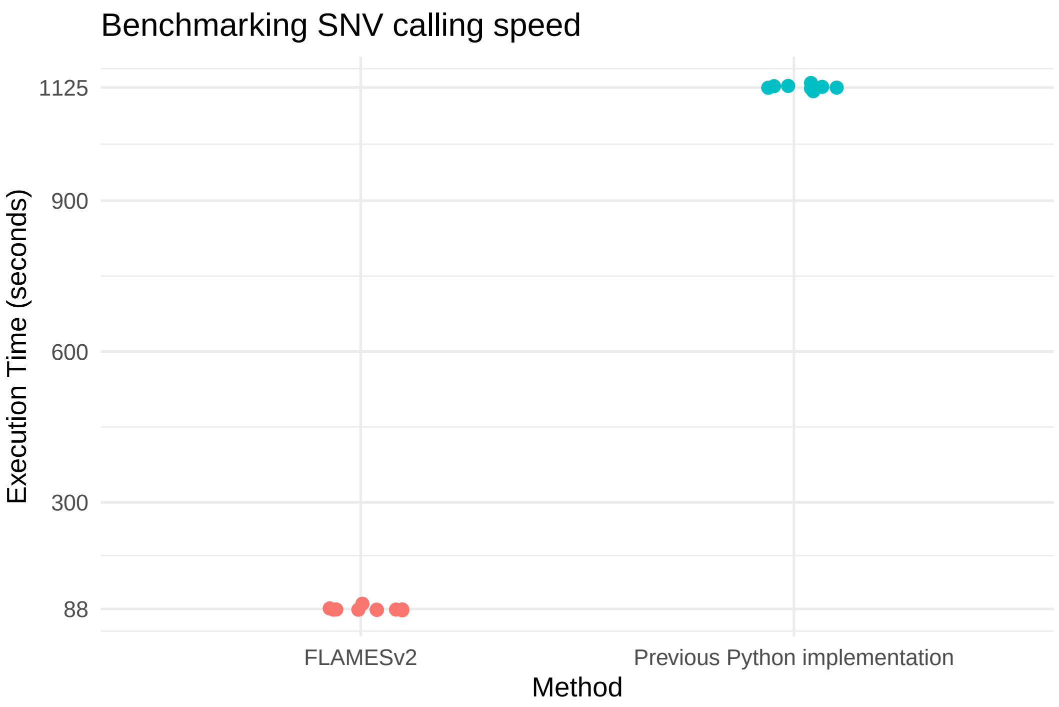


**Fig. S2. Single-cell SNV quantification speed comparison of `FLAMESv2` with previous Python implementation.** Compared using a $\sim$ 1 GiB `BAM` file filtered to keep alignments covering 5 loci, running both version of `FLAMES` 10 times with 4 cores and 16 GiB of RAM.

| **Metric** | **`FLAMESv1`** | **`FLAMESv2`** |
| --- | --- | --- |
| Cells identified | 337 | 339 |
| Number of unique genes | 5,578 | 29,013 |
| Number of isoforms | 15,430 | 93,721 |
| Median genes per cell | 2,390 | 7,182 |
| Median isoforms per cell | 3,334 | 10,812 |
| Median reads per cell | 12,922 | 31,923 |
| Number of novel isoforms* | 5,834 | 2,855 |
| Incomplete transcripts (ISM) | 236 | 0 |
| Usable reads (reads assigned to a cell) | 71% | 77% |
| Reads used to quantify isoforms** | 5,135,544 | 14,586,085 |

**Table S2. Comparison of FLAMES versions across key metrics.** Genes identified include protein coding and non-coding. * Novel isoforms not classified as FSM or ISM by SQANTI3. ** Summed counts in the isoform counts file. Reads used to quantify isoforms have been demultiplexed and deduplicated. Novel Bambu genes have been excluded from the `FLAMESv2` metrics.

| Sample | Number of pass reads | Percentage of usable reads | Saturation | Number of cells identified by FLAMESv2 | Number of high quality cells | Median gene counts | Median isoforms counts | Median features gene | Median features isoforms |
|---|---|---|---|---|---|---|---|---|---|
| C1-STC | 70,630,939 | 91.26 | 12.93 | 1195 | 849 | 44648 | 35598 | 8890 | 12998 |
| C4-STC | 62,545,336 | 89.76 | 32 | 1302 | 997 | 29076 | 20278 | 7681 | 10940 |
| C2-Day25 | 69,190,113 | 90.21 | 39.94 | 828 | 665 | 35082 | 23642 | 7925 | 10883 |
| C5-Day25 | 169,295,516 | 86.07 | 56.02 | 2843 | 1040 | 35623 | 26312 | 7534 | 9626 |
| C2-Day55 | 158,640,053 | 81.42 | 41.8 | 1876 | 1668 | 19490 | 11812 | 6520 | 8239 |
| C3-Day55 | 147,964,083 | 82.32 | 38.66 | 1470 | 1330 | 39018 | 20037 | 8619 | 11105 |
| C3-Day80 | 128,759,089 | 81.93 | 30.43 | 1199 | 1048 | 46631 | 23449 | 9714 | 12996 |
| C5-Day80 | 142,263,891 | 86.67 | 24.7 | 2245 | 2051 | 28675 | 17308 | 7786 | 10359 |

**Table S3. LR scRNA-seq summary metrics.** Pass reads have a Qscore > 10. Usable reads are those that have the correct read structure, i.e: a cell barcode, UMI, and unambiguous polyT and adapter positions found by BLAZE.

| Contrast | DTUrtle | IsoformSwitchAnalyzer | Number of overlaps | Union | Jaccard |
|---|---|---|---|---|---|
| Day80 vs STC | 8850 | 10539 | 5797 | 13592 | 0.4265009 |
| Day55 vs STC | 8266 | 9920 | 5332 | 12854 | 0.4148125 |
| Day55 vs Day80 | 2541 | 636 | 358 | 2819 | 0.1269954 |
| Day25 vs STC | 5472 | 3923 | 2115 | 7280 | 0.290522 |
| Day25 vs Day80 | 7623 | 8365 | 4426 | 11562 | 0.3828057 |
| Day25 vs Day55 | 5906 | 5889 | 3064 | 8731 | 0.3509335 |
| Overall | 15794 | 18490 | 11163 | 23121 | 0.4828078 |

**Table S4.** Concordance of Differential Transcript Usage (DTU) between DTUrtle and IsoformSwitchAnalyzer across time-point contrasts. For each contrast, we report per-tool DTU counts, their intersection (Number of overlaps), Union and Jaccard similarity (overlap/union). “Overall” aggregates unique DTU events across all contrasts to estimate global agreement.

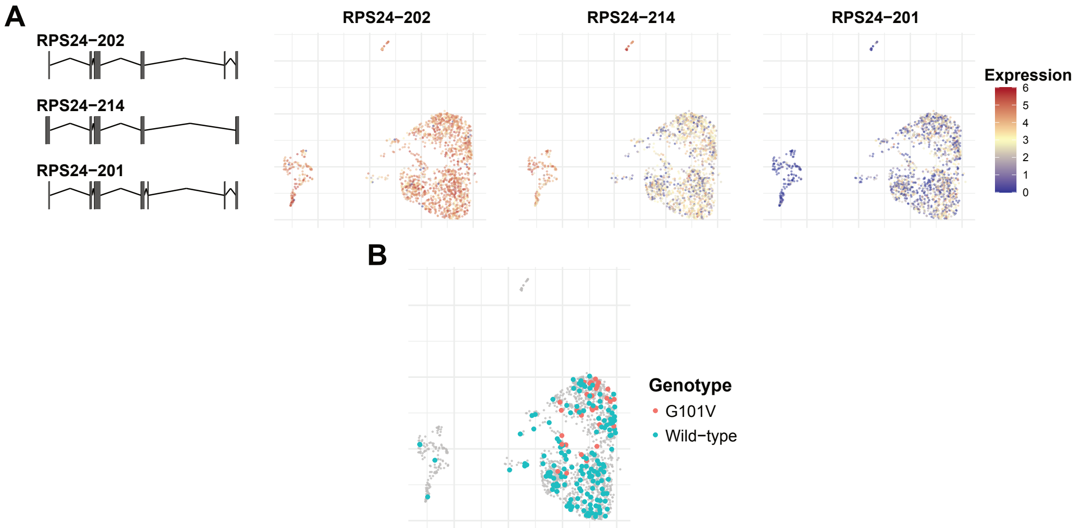


**Fig. S3. Analysis of peripheral blood mononuclear cells (PBMCs) from a chronic lymphocytic leukemia (CLL) patient** (**A**) UMAPs of CLL2 data from Tian *et al.* showing expression of *RPS24* isoforms created with `FLAMESv2` using UMAP embeddings from Tian *et al.* [2] (**B**) UMAP of CLL2 data coloured by *BCL2 G101V* allele created with `FLAMESv2` using UMAP embeddings from Tian *et al.*

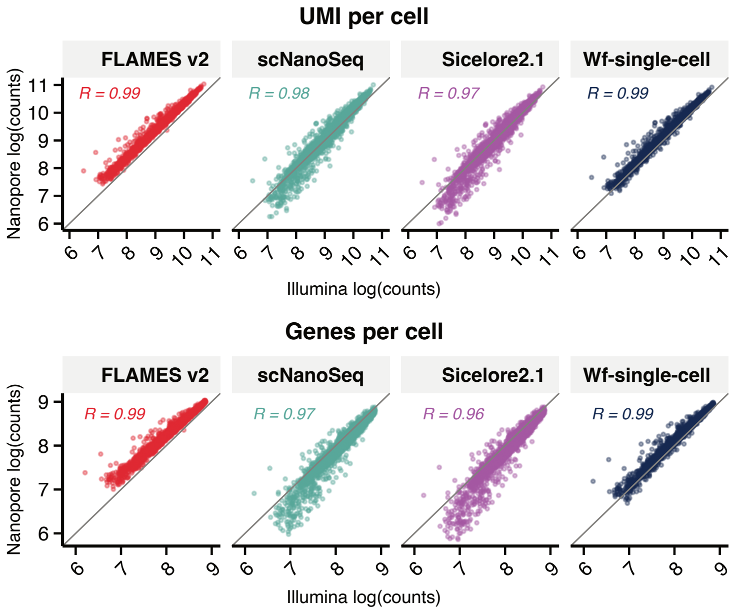


**Fig. S4. Per-cell UMI and gene detection concordance between long-read methods and Illumina short-read sequencing.** Scatter plots comparing per-cell total UMI counts (top) and number of detected genes (bottom) between Illumina (x-axis) and four long-read single-cell methods (y-axis). Values are log-transformed (log1p). The diagonal grey line indicates perfect concordance. Pearson correlation coefficients (R) are shown for each method. Analysis was performed on a shared gene and cell universe to ensure direct comparability across method (19424 genes across 3942)

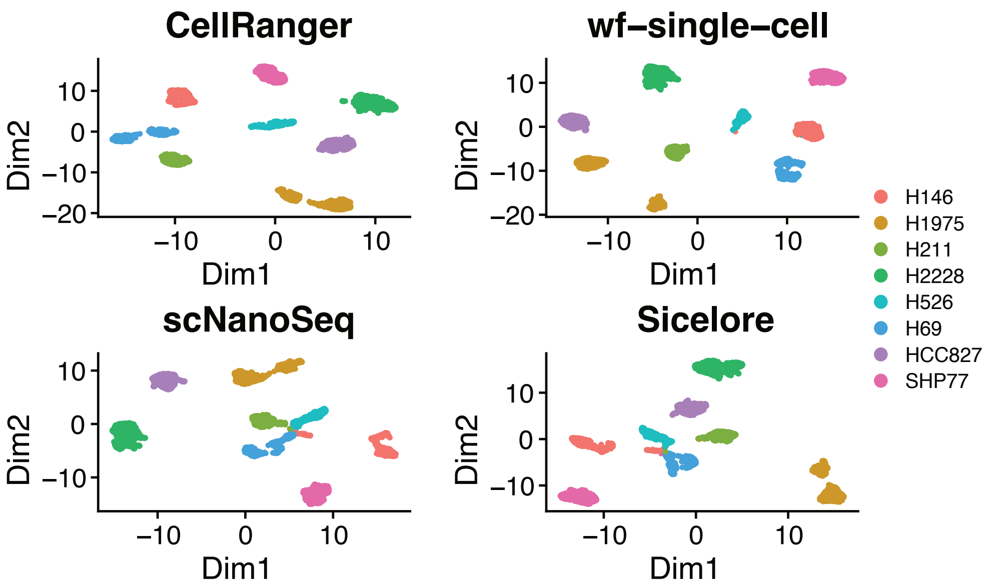


**Fig. S5. UMAP visualization of LongBench ONT single-cell dataset.** UMAP visualization generated from CellRanger (short-read), wf-single-cell (long-read), scNanoSeq (long-read) and SiCeLoRe (long-read) results on the LongBench single-cell datasets, with cell lines colored according to SNP-based annotation.

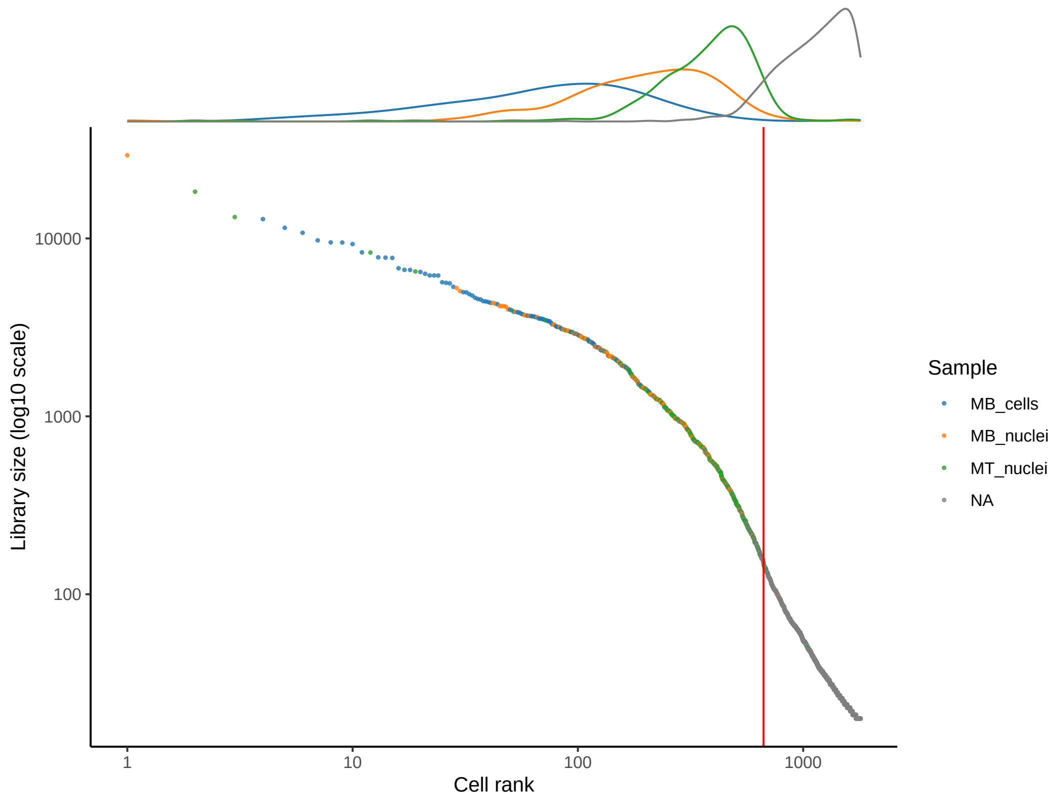


**Fig. S6. Barcode knee-plot for LR-split-seq data from Rebboah *et al.* [3] produced by `FLAMESv2`.** UMI counts from `FLAMESv2`'s gene count matrix were used to compute library size and cell-type annotation from Rebboah *et al.*'s filtered short-read cells were used to color cells, with cells not present in filtered short-read cells in grey. Density plot of the annotation were shown on top.

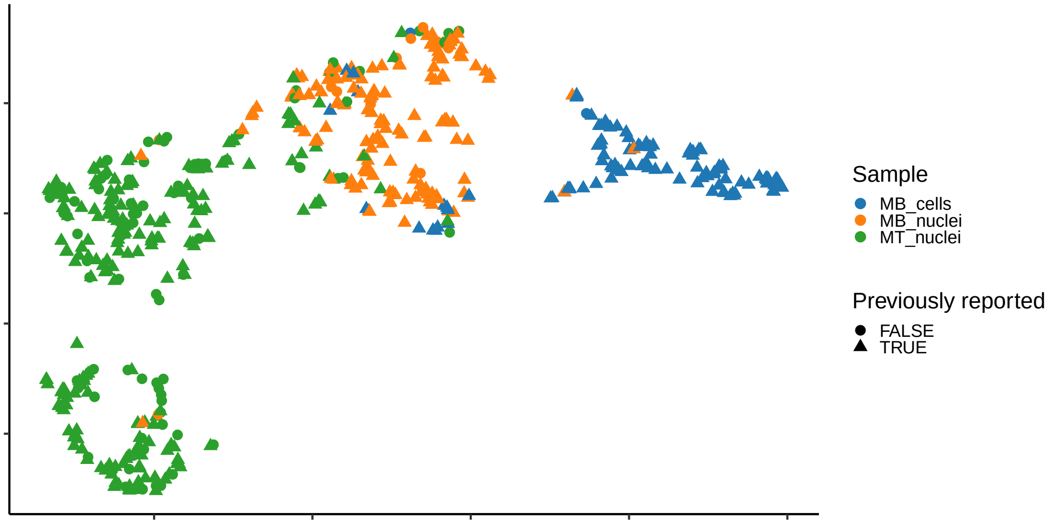


**Fig. S7. UMAP of LR-split-seq data derived using gene counts from *FLAMESv2* using filtered barcodes as demultipex allow-list [3]**

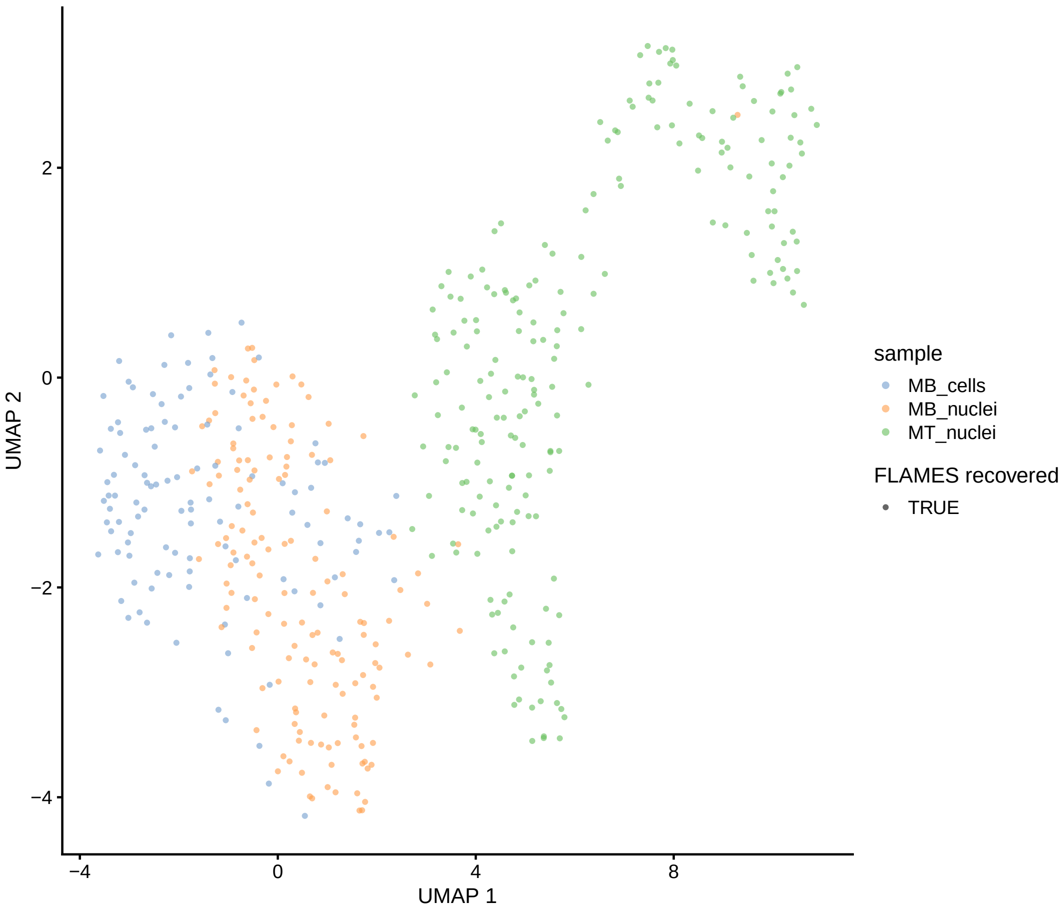


**Fig. S8. UMAP derived using gene counts from *LR-splitpipe* from Rebboah *et al.* [3], coloured by sample.**

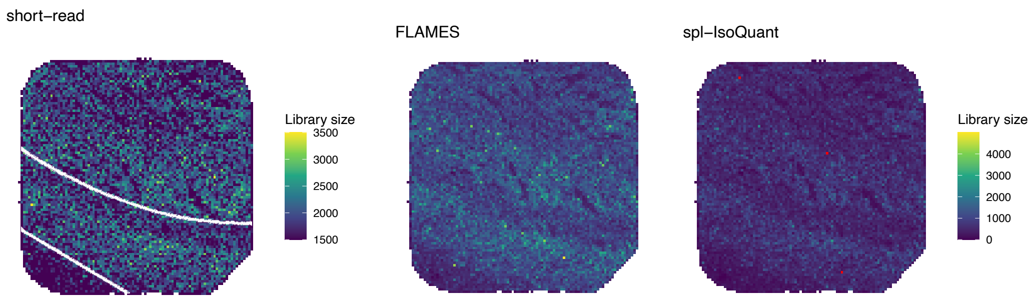


**Fig. S9. Library size of the Curio sample for the coupled short-read processed by Curio Seeker and long-reads processed by *FLAMESv2* and *spl-IsoQuant* respectively**. The spatial spots were aggregated to 50 µm bins.

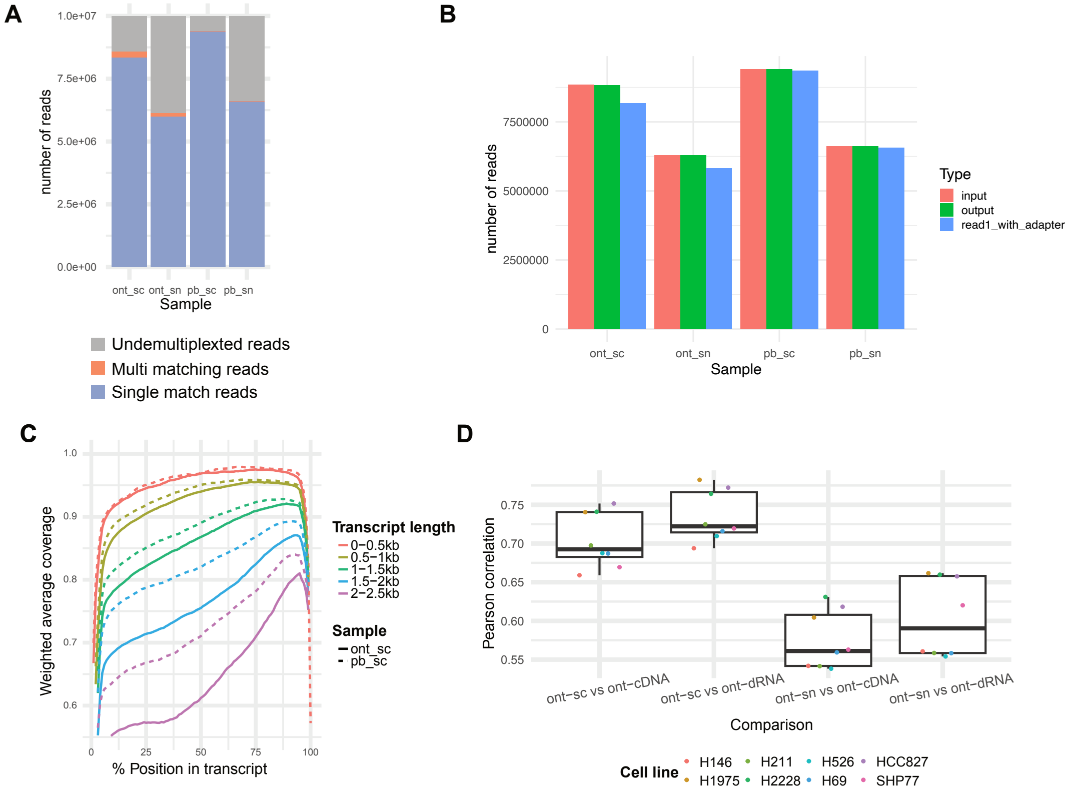


**Fig. S10. Benchmarking different long-read sequencing protocols with `FLAMESv2`**

(**A**) `FLAMESv2` demultiplexing results for LongBench [1] single-cell (sc) and single-nuclei (sn) with Nanopore (ont) and PacBio (pb) sequencing samples 10 million reads were used for each sample. (**B**) *cutadapt* result of LongBench single-cell and single-nuclei with Nanopore and PacBio sequencing samples. `read1_with_adapter` indicates reads where TSO sequence were found and trimmed. Single-cell (sc), single-nuclei (sn), Nanopore (ont), PacBio (pb). (**C**) Average sequencing coverage along the transcript body of LongBench Nanopore and PacBio sequencing samples. (**D**) Pearson correlation of gene expression counts between ONT bulk (direct RNA, PCR-cDNA) and ONT pseudobulk (single nucleus and single cell) data for each cell line in the LongBench data.

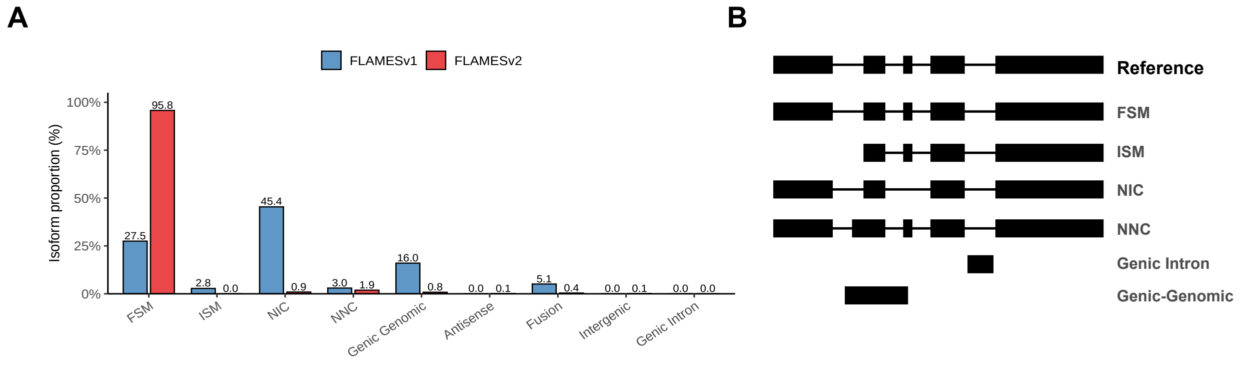


**Fig. S11.** **SQANTI comparison of FLAMESv1 and FLAMESv2 isoform calls.** **(A)** SQANTI structural categories of isoforms identified by FLAMESv1 and FLAMESv2 on identical data. Isoform discovery in FLAMESv1 was performed using the original implementation, whereas FLAMESv2 employs Bambu for novel isoform discovery. All novel Bambu genes were removed prior to running SQANTI3. **(B)** Schematic representation of SQANTI structural categories, adapted from [4, 5]. FSM - Full splice match, ISM - Incomplete splice match, NIC - Novel in catalogue, NNC - Novel not in catalogue.

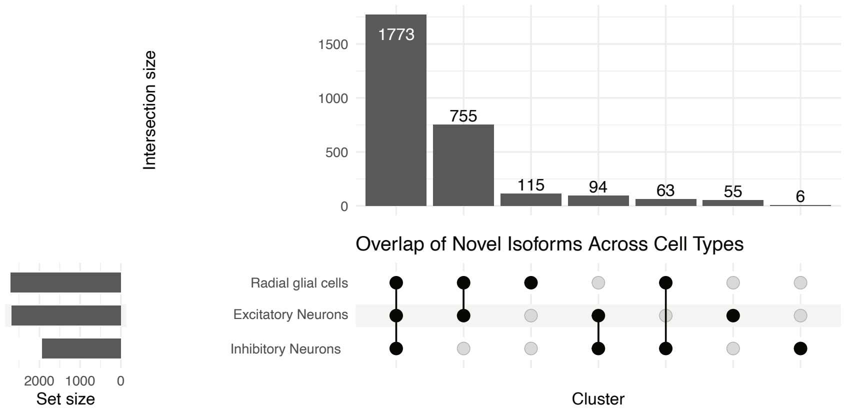


**Fig. S12. UpSet plot showing the overlap of novel isoforms across radial glia, excitatory neurons, and inhibitory neurons.** Each bar on the left indicates the total number of novel isoforms detected in a given cell type, and the intersection matrix on the bottom highlights how many of those isoforms are uniquely expressed or shared between cell types.

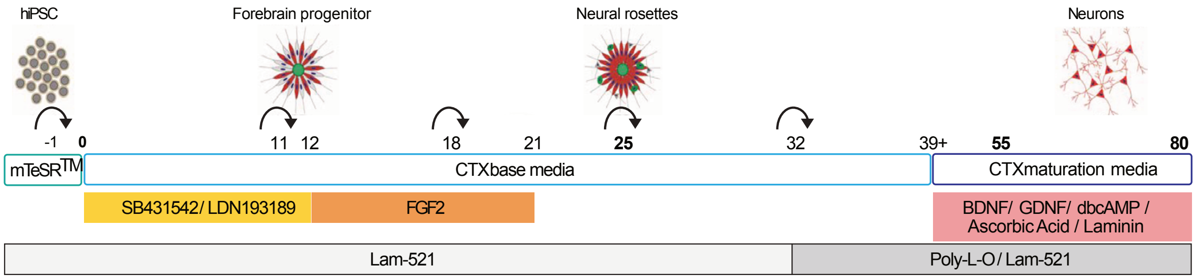


**Fig. S13. hiPSC to neuron differentiation**. Coloured boxes represent the growth factors that the differentiation is exposed to. Rounded arrows indicate a cell passage and bold numbers indicate sample collection time for disassociation and single-cell preparation. We collected samples along the 80 day differentiation at the following developmental stages: iPSC – beginning of the differentiation, D25 - neural progenitor expansion and the formation of neural rosettes, D55 and Day80 – neurogenesis. Day -1, refers to day prior to neural induction.

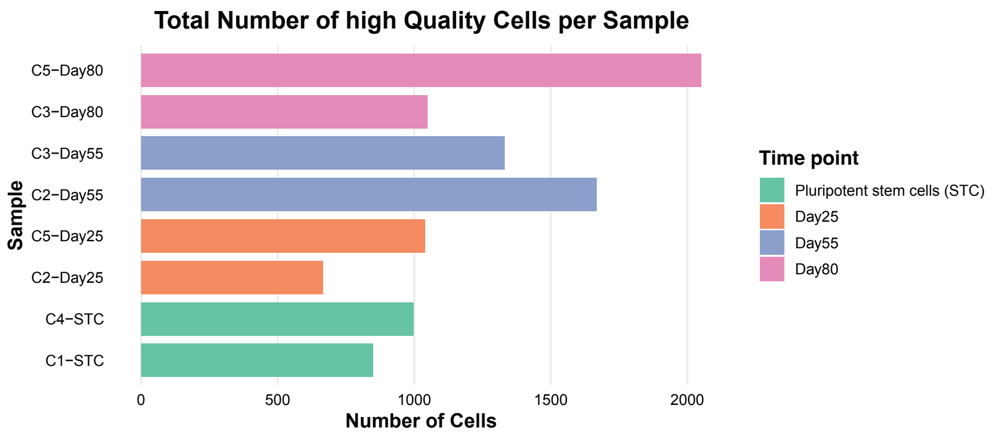


**Fig. S14. High-quality cells passing QC.** Bar plots show the number of high-quality cells that passed quality control across samples. Sample names use the **C** prefix to indicate the collection window, such that samples with the same **C** number were collected at the same time. STC is used to refer to pluripotent stem cells

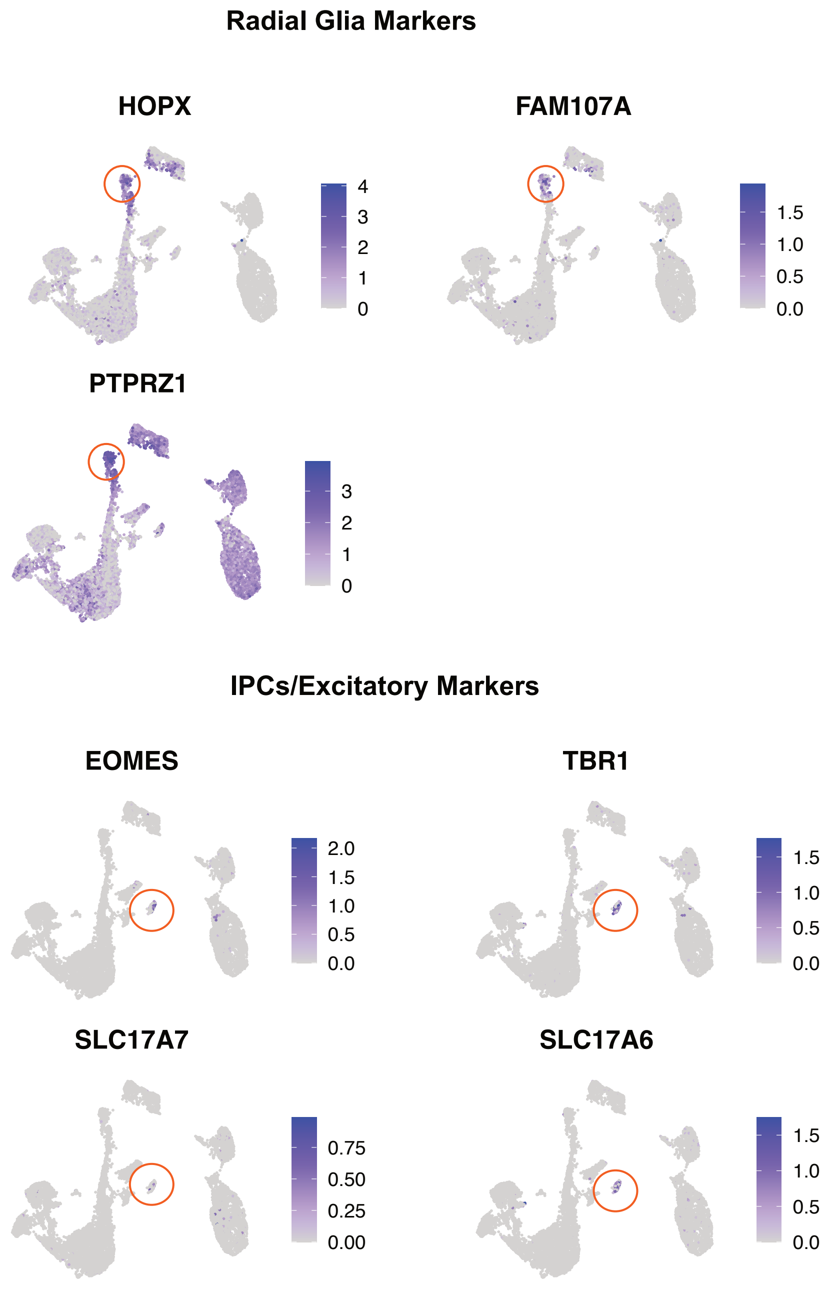


**Fig. S15. UMAP expression of marker genes for key cell types.** Radial glia are identified by expression of *HOPX*, *FAM107A*, and *PTPRZ1* (top row). Intermediate progenitors (IPCs) / Excitatory neurons are identified by expression of *EOMES*, *TBR1*, *SLC17A7*, and *SLC17A6* (bottom row). Red circles highlight representative clusters enriched for these markers.

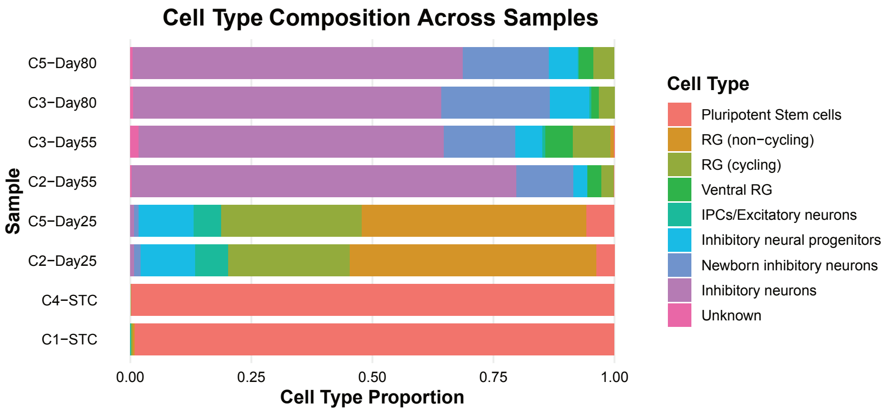


**Fig. S16. Cell type composition across samples.** Bar plots show the proportional representation of major cell types identified at each developmental stage and sample. Sample names use the **C** prefix to indicate the collection window, such that samples with the same **C** number were collected at the same time.

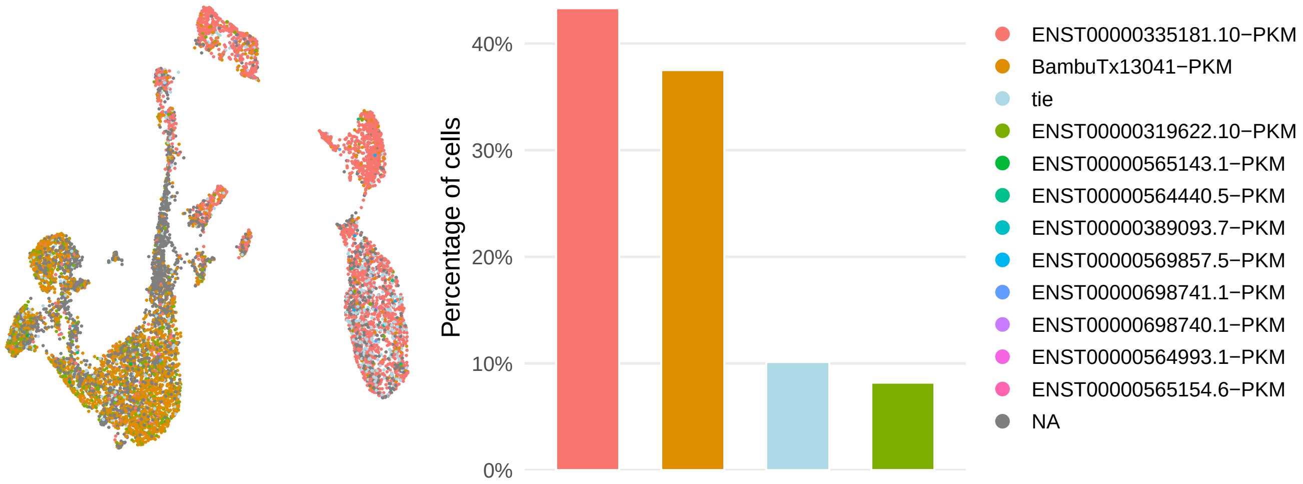


**Fig. S17. *PKM* isoform switching.** Left: UMAP visualization of cells coloured by the dominant *PKM* isoform expressed in each cell. The dominant isoform was defined as the transcript with the highest count within that cell. Grey (NA) denotes cells with less than 3 *PKM* counts. Right: Bar plot showing the percentage of cells in which each isoform is dominant. Tie denotes cells with two or more dominant *PKM* isoforms

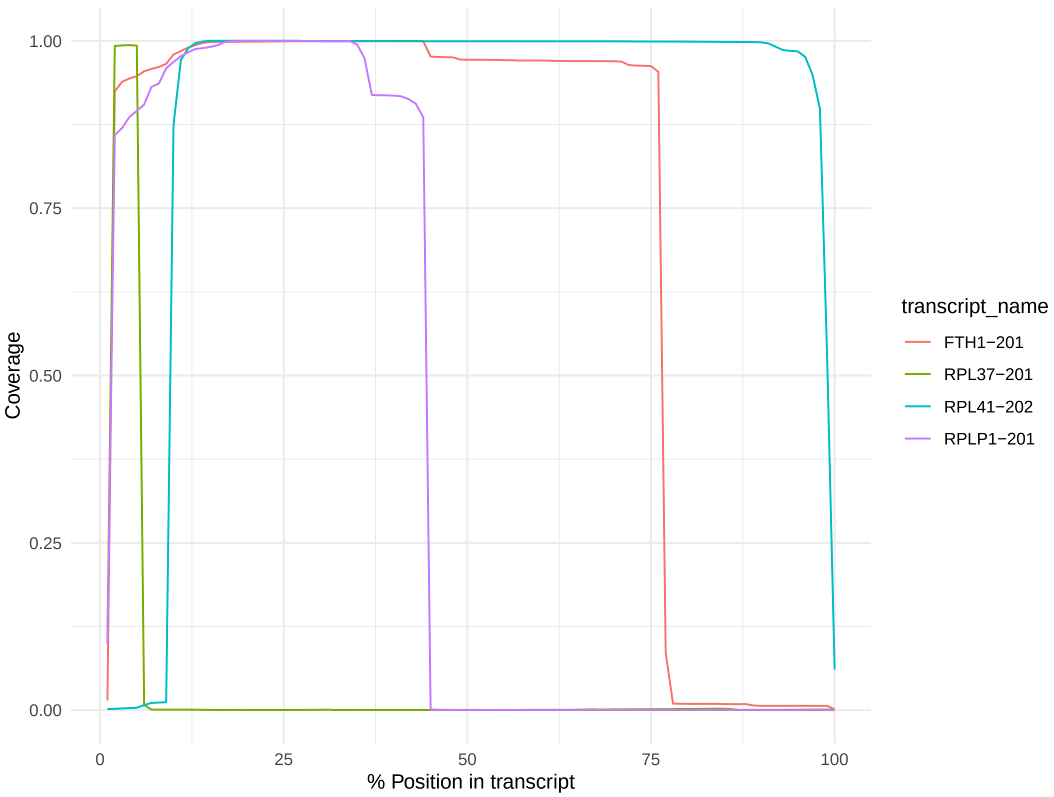


**Fig. S18. Coverage plot of filtered transcripts.** Top 4 transcripts by read counts removed by `FLAMESv2`'s `filter_coverage` function due to having drastic coverage changes in the Nanopore single-cell sample from LongBench. Coverage (number of reads scaled by total number of aligned reads) shown on y-axis, sampling at every percentile of the transcript length from 5' to 3' and showing the percentiles on the x-axis.

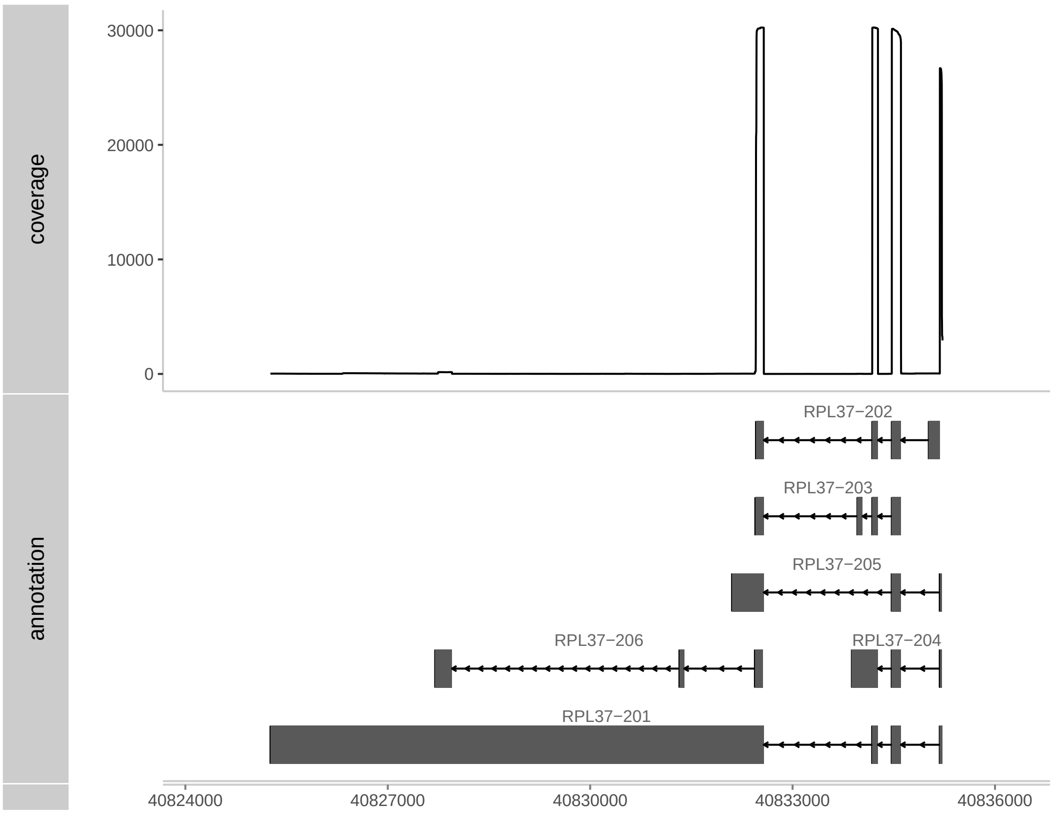


**Fig. S19. Coverage and gene annotation of *RPL37* gene.** Genomic coverage of the *RPL37* gene (top) and its reference annotations (bottom), highlighting the potential error in annotation causing *RPL37-201* (shown previously in Figure S18) to be filtered.
